# Lysosomal TBK1 Responds to Amino Acid Availability to Relieve Rab7-Dependent mTORC1 Inhibition

**DOI:** 10.1101/2023.12.16.571979

**Authors:** Gabriel Talaia, Amanda Bentley-DeSousa, Shawn M. Ferguson

## Abstract

Lysosomes play a pivotal role in coordinating macromolecule degradation and regulating cell growth and metabolism. Despite substantial progress in identifying lysosomal signaling proteins, understanding the pathways that synchronize lysosome functions with changing cellular demands remains incomplete. This study uncovers a role for TANK-binding kinase 1 (TBK1), well known for its role in innate immunity and organelle quality control, in modulating lysosomal responsiveness to nutrients. Specifically, we identify a pool of TBK1 that is recruited to lysosomes in response to elevated amino acid levels. At lysosomes, this TBK1 phosphorylates Rab7 on serine 72. This is critical for alleviating Rab7-mediated inhibition of amino acid-dependent mTORC1 activation. Furthermore, a TBK1 mutant (E696K) associated with amyotrophic lateral sclerosis and frontotemporal dementia constitutively accumulates at lysosomes, resulting in elevated Rab7 phosphorylation and increased mTORC1 activation. This data establishes the lysosome as a site of amino acid regulated TBK1 signaling that is crucial for efficient mTORC1 activation. This lysosomal pool of TBK1 has broader implications for lysosome homeostasis, and its dysregulation could contribute to the pathogenesis of ALS-FTD.

## Introduction

In addition to degrading macromolecules that are delivered by the endocytic and autophagy pathways, lysosomes also act as signaling platforms that control cell growth and metabolism (Ballabio and Bonifacino, 2019; Ferguson, 2015; Goul et al., 2023). Coordinating the degradative and signaling functions of lysosomes is crucial because it ensures that cells can efficiently recycle damaged or surplus components while precisely regulating their growth and energy use according to the cell’s needs and external cues. Mechanistic target of rapamycin complex 1 (mTORC1) is a kinase that communicates between lysosomes and the rest of the cell to help match growth to the availability of nutrients and growth factors (Goul et al., 2023; Liu and Sabatini, 2020). To this end, multiple nutrients are sensed and integrated upstream of the heterodimeric Rag GTPases that recruit mTORC1 to the surface of lysosomes (Goul et al., 2023; Lama-Sherpa et al., 2023; Liu and Sabatini, 2020). Meanwhile, growth factor signaling controls the Rheb GTPase which is responsible for mTORC1 activation at lysosomes (Angarola and Ferguson, 2020; Menon et al., 2014). This complex coordination between diverse inputs to regulate signals from lysosomes highlights the critical need for cells to balance their degradative activities with overall nutrient requirements.

Cells must also monitor and manage lysosome integrity to ensure that lysosomes remain intact and lysosomal enzymes do not leak into the cytoplasm (Bohannon and Hanson, 2020; Yang and Tan, 2023). When lysosomes suffer severe damage that cannot be repaired, TANK-binding kinase 1 (TBK1) helps cells to clear severely damaged lysosomes via the process known as lysophagy where the damaged lysosomes are engulfed by autophagosomes and then delivered to intact lysosomes for degradation (Eapen et al., 2021; Hung et al., 2013; Maejima et al., 2013; Pied et al., 2022; Vargas et al., 2023). TBK1 also plays important roles in the autophagic clearance of damaged mitochondria and in signaling related to innate immunity (Fitzgerald et al., 2003; Harding et al., 2021; Heo et al., 2018; Moore and Holzbaur, 2016; Pomerantz and Baltimore, 1999). Heterozygous loss-of-function mutations in the *TBK1* gene cause amyotrophic lateral sclerosis (ALS) and frontotemporal dementia (FTD) (Cirulli et al., 2015; Freischmidt et al., 2015; Gijselinck et al., 2015). TBK1 missense mutations have also been identified as causes of ALS and FTD (Freischmidt et al., 2015; Gijselinck et al., 2015; Hirsch-Reinshagen et al., 2019; Van Mossevelde et al., 2016). Heterozygous loss-of-function mutations in TBK1 also cause herpes simplex encephalitis (Ahmad et al., 2016). Meanwhile, excess TBK1 activity arising from gene duplication causes open angle glaucoma (Ahmad et al., 2016). Aberrant TBK1 activity can also contribute to diabetes and obesity (Bodur et al., 2022; Cruz et al., 2018; Oral et al., 2017; Reilly et al., 2013). These diseases arising from both too little and too much TBK1 activity demonstrate the importance of regulatory mechanisms that ensure tight control of TBK1 kinase activity. This breadth of physiological and pathophysiological functions for TBK1 also raises questions about underlying subcellular sites of action and relevant substrates.

In addition to substrates directly related to innate immunity, TBK1 phosphorylates mTORC1 components leading to both activation and inhibition of mTORC1 kinase activity in different studies (Antonia et al., 2019; Bodur et al., 2018; Cooper et al., 2017; Hasan et al., 2017; Ye et al., 2023). These reports of opposing effects of TBK1 on mTORC1 activity are puzzling and raise questions about underlying mechanisms. As mTORC1 activation is now well known to take place at lysosomes, if TBK1 directly regulates mTORC1 or its immediate upstream regulators, then TBK1 should also be present on intact lysosomes and not just recruited in response to severe damage that induces lysophagy (Eapen et al., 2021; Goul et al., 2023). If so, there should also be signals that control TBK1 at intact lysosomes and lysosome-localized TBK1 substrates that mediate communication from TBK1 to mTORC1.

To investigate lysosomal functions of TBK1 and to define the relationship between TBK1 and mTORC1 activation, we employed a series of genetic and pharmacological perturbations in mammalian cells in culture combined with assays of TBK1 subcellular localization, substrate phosphorylation and mTORC1 activity. We observed that both acute pharmacological inhibition and genetic depletion of TBK1 resulted in a significant reduction in the ability of mTORC1 to be efficiently activated in response to feeding cells with amino acids. This need for TBK1 in mTORC1 activation was paralleled by observations that TBK1 is recruited to the surface of lysosomes concurrent with mTORC1 activation when starved cells are re-fed with amino acids. We furthermore established that TBK1-dependent phosphorylation of Rab7 (serine 72) is critical for relieving inhibition of mTORC1 by Rab7. Finally, we discovered that the ALS-FTD TBK1-E696K mutant is constitutively localized to lysosomes, more active and more strongly promotes mTORC1 activation by amino acids. Altogether, this research demonstrates that amino acid availability controls the ability of a lysosomal pool of TBK1 to relieve Rab7-dependent inhibition of mTORC1 activation.

## Results

### TBK1 promotes amino acid-dependent activation of mTORC1 at the lysosomes

To test for TBK1 functions upstream of mTORC1 at lysosomes, we performed genome editing to knock out TBK1 in HeLa cells and measured the impact on mTORC1 activation in assays where starved cells were re-fed with amino acids. These experiments revealed reduced mTORC1 activation by amino acids in the TBK1 KO cells as measured by phosphorylation of its direct substrates, S6K1 (phospho-T389) and ULK1 (phospho-S757; Fig. 1A and B, S1A). These two proteins are well-established mTORC1 substrates whose phosphorylation is regulated by amino acid availability (Liu and Sabatini, 2020). Similar results were obtained from mouse RAW264.7 macrophage cells when we knocked out both TBK1 and the closely related IKKε homolog that is expressed in myeloid cells (Fig. S1B and C)(Balka et al., 2020). To rule out long term adaptation to the TBK1 KO as a contributing factor to our observations, we performed acute treatment of RAW264.7 cells with BX-795, an inhibitor of TBK1 kinase activity, and this yielded a similar effect to TBK1 KO on mTORC1 activation (Fig. 1C and D)(Bain et al., 2007). Furthermore, the mTORC1 activation defect in TBK1 KO HeLa cells was fully rescued following stable expression of TBK1- GFP (Fig. 1E and F). Confocal immunofluorescence microscopy revealed the presence of TBK1 puncta adjacent to lysosomes in wild type HeLa cells and this was lost in the TBK1 KO cells. In the TBK1 KO cells that were rescued with TBK1-GFP at ∼4x above endogenous levels, the lysosome localization of TBK1 was more evident (Fig. 1G). TBK1 lysosome localization was independently assessed by immunoblotting of lysosomes that were magnetically isolated from cells whose lysosomes were endocytically loaded with superparamagnetic iron oxide nanoparticles (SPIONs) via a pulse-chase strategy (Amick et al., 2018; Rodriguez-Paris et al., 1993; Tharkeshwar et al., 2020). Both endogenous TBK1 and TBK1-GFP were present on the purified lysosomes along with established late endosome/lysosome proteins (LAMP1 and Rab7) (Fig. 1H). As expected, PDI and GM130, proteins of the endoplasmic reticulum and Golgi, were depleted from the purified lysosome samples. Rab7, a small GTPase that regulates multiple aspects of late endosomes and lysosomes, was also abundant in the purified lysosomes samples (Kummel et al., 2023). Rab7 is a major substrate of TBK1 but the functional significance of this has not been extensively studied (Heo et al., 2018; Ritter et al., 2020). Interestingly, Rab7 phosphorylated on serine 72 (S72) was detected in both the whole cell lysates and purified lysosomes. The abundance of this phospho-Rab7 was reduced in the TBK1 KO cells and increased in the TBK1-GFP rescue line (Fig. 1H). To ensure that this TBK1-GFP fusion protein supports normal TBK1 regulation and functions, we treated TBK1 KO cells that express TBK1- GFP with cGAMP [an agonist for Stimulator of Interferon Genes (STING), a robust activator of TBK1] and confirmed that this resulted in the expected TBK1 autophosphorylation on S172, an interaction between TBK1 and STING and a band shift for STING that reflects its phosphorylation downstream of TBK1 (Fig. S1D) (Tanaka and Chen, 2012).

**Figure 1:**
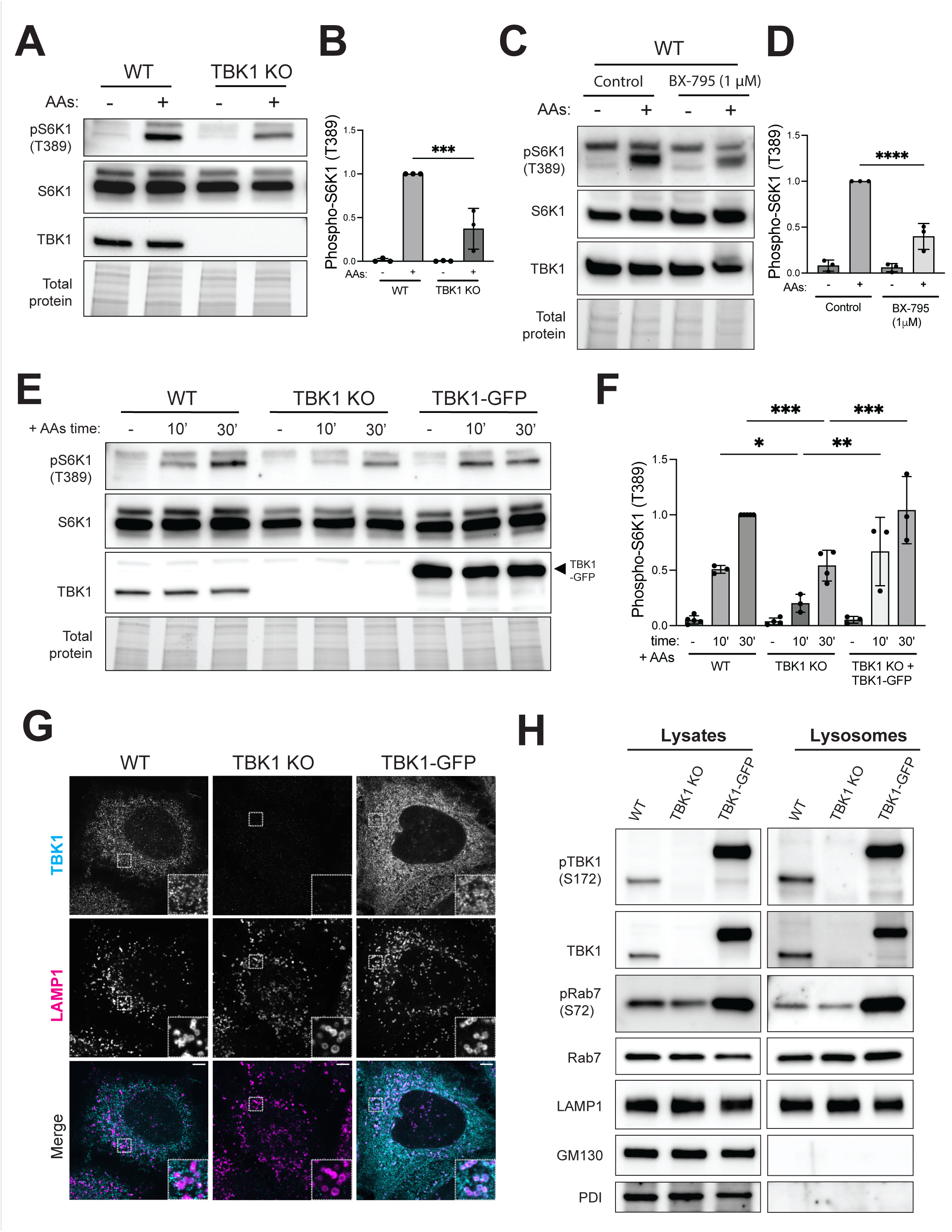
Lysosomal TBK1 promotes amino acid-dependent mTORC1 signaling. (A) Immunoblot analysis of S6K1 phosphorylated at Thr389 [pS6K1 (T389)], total S6K1 and TBK1 in whole-cell lysates from WT versus TBK1 KO HeLa cells that were starved of amino acids for 60 minutes (-) and then re-fed with amino acids for 60 minutes (+). TGX stain-free method was used to visualize total protein. (B) Quantification of phospho-S6K1 (T389) normalized to total S6K1 and expressed as a fold change compared to the WT cells under re-fed conditions. Statistical significance was determined by ordinary one-way analysis of variance (ANOVA) with Šidák *post hoc* test (n=3; mean ± SD; ***, *p*<0.001). (C) Immunoblot analysis of RAW264.7 cells that underwent amino acid starvation (60 minutes) and re-feeding (60 minutes) in the presence of 1 µM BX-795 or 0.01% (v/v) DMSO (Control). (D) Quantification of phospho-S6K1 (T389) normalized to total S6K1 and expressed as a fold change compared to the WT cells under re-fed conditions. Statistical significance was determined by ordinary one-way analysis of variance (ANOVA) with Šidák *post hoc* (n=3; mean ± SD; ****, *p*<0.0001). (E) Immunoblots from wildtype (WT), TBK1 KO and TBK1 KO stably expressing TBK1-GFP (TBK1-GFP) HeLa cells that were starved for 60 min (-) and then re-fed with amino acids for 10 or 30 minutes (10’ and 30’). TGX stain-free method was used as a loading control (Total protein). (F) Phospho-S6K1 levels were quantified and normalized to total S6K1 from at least three independent experiments, expressed as fold change compared to the WT cells under 30 minutes re-fed conditions. Statistical significance was determined by ordinary one-way analysis of variance (ANOVA) with Šidák *post hoc* (mean ± SD; ***, *p*<0.001; **, *p*<0.01; *, *p*<0.05). (G) High resolution immunofluorescence microscopy of TBK1 and LAMP1 in wildtype, TBK1 KO and TBK1-GFP HeLa cells under basal growth conditions. Insets are magnified 3X. Scale bar: 5 µm. (H) Immunoblot analysis of the indicated proteins in cell lysates (Lysates) and magnetically isolated lysosomes (Lysosomes) from WT, TBK1 KO and TBK1-GFP HeLa cells under basal growth conditions. Blots for Lysate and Lysosome fractions were obtained in parallel from the same membrane.

### TBK1 signaling at lysosomes is stimulated when cells are fed with amino acids

TBK1-dependent phosphorylation of Rab7 under basal cell growth conditions was quantified in whole cell lysates from wild type, TBK1 KO and TBK1-GFP rescued HeLa cells. Rab7 phosphorylation was decreased in the TBK1 KO cells and was rescued to above wild type levels following the stable over-expression of TBK1-GFP in the TBK1 KO background (Fig. 2A and B). Phosphorylation of Rab7 by TBK1 was previously described to occur during the very specific contexts of mitophagy and innate immunity signaling in the STING pathway (Heo et al., 2018; Ritter et al., 2020). Our observations of TBK1 at lysosomes with impacts on Rab7 phosphorylation and mTORC1 activation in the absence of stimuli related to mitochondrial damage or STING activation raised two questions. First, what are the inputs that regulate TBK1-dependent control of mTORC1 activity? Second, does Rab7 phosphorylation contribute to TBK1-dependent activation of mTORC1? Answering these questions was important for establishing a functional relevance for TBK1 at lysosomes.

**Figure 2:**
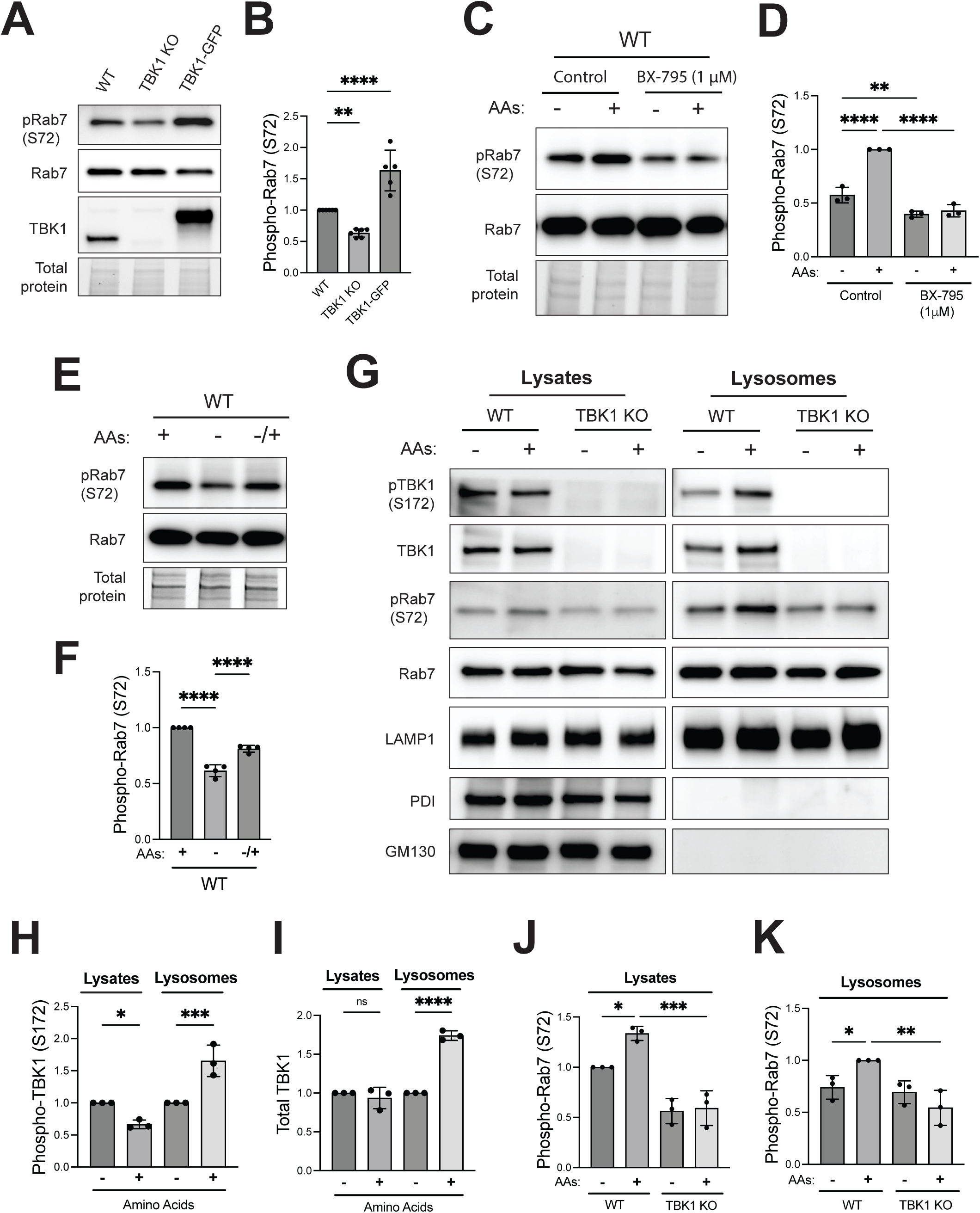
TBK1-dependent Rab7 phosphorylation is regulated by amino acid availability. (A) Immunoblot analysis of WT, TBK1 KO and TBK1 KO + TBK1-GFP HeLa cells under basal conditions. (B) Phospho-Rab7 (S72) protein levels from panel A were quantified and normalized to total Rab7 (WT is considered 1). Data of at least five independent experiments was plotted and statistical significance was determined by ordinary one-way ANOVA with Šidák *post hoc (*mean ± SD; ****, *p*<0.0001; **, *p* <0.01). (C) Immunoblot analysis of RAW264.7 cells that underwent amino acid starvation (60 minutes) and re-feeding (60 minutes) in the presence of 1 µM BX-795 or 0.01% DMSO (Control). (D) Phospho-Rab7 protein levels were quantified and normalized to total Rab7 (re-fed or untreated WT is considered 1). Statistical significance was determined by ordinary one-way ANOVA with Šidák post-test (n=3; mean ± SD; ****, *p*<0.0001; **, *p* <0.01). (E) Immunoblot analysis of pRab7 (S72), total Rab7 in whole-cell lysates of WT HeLa cells under basal conditions (+), starved (-) and amino acid re-fed (-/+). (F) Phospho-Rab7 (S72) protein levels were quantified and normalized to total Rab7 (basal conditions were considered 1). Statistical significance was determined by ordinary one-way ANOVA with Šidák post-test (n=4; mean ± SD; ****, *p*<0.0001). Total protein was detected by the TGX stain-free method. (G) Immunoblot analysis of the indicated proteins in cell lysates (Lysates) and magnetically isolated lysosomes (Lysosomes) of WT and TBK1 KO HeLa cells starved for 60 min (-) and re-fed with amino acids for 60 min (+). Blots for Lysate and Lysosome fractions were obtained in parallel from the same membrane. (H-I) Quantification of phosphorylated TBK1 (S172) and total TBK1 normalized to total Rab7 in Lysates and Lysosomes, expressed as fold change compared to starved cells. (J-K) Phospho-Rab7 (S72) was quantified and normalized to total Rab7 in Lysates and Lysosomes. Data points from three independent experiments and statistical significance was determined by ordinary one-way ANOVA with Šidák *post hoc* (mean ± SD; ****, *p*<0.0001; ***, *p*<0.001; **, *p*<0.01; *, *p*<0.05; ns, not significant).

Given that TBK1 is required for efficient mTORC1 activation in response to the addition of amino acids, we tested whether TBK1 activity is also regulated by this stimulus. Indeed, Rab7 phosphorylation increased when starved RAW264.7 cells were re-fed with amino acids and both basal and amino acid-stimulated Rab7 phosphorylation was blocked by acute treatment with BX- 795, an inhibitor of TBK1 kinase activity (Fig. 2C and D). HeLa cells also showed a drop in Rab7 S72 phosphorylation in response to starvation and restoration of Rab7 phosphorylation in response to amino acid re-feeding (Fig. 2 E and F). In support of the functionality of the TBK1- GFP chimeric protein, TBK1 KO HeLa cells that were rescued with TBK1-GFP also showed a similar pattern of reduced Rab7 phosphorylation upon starvation and an increase in response to amino acid re-feeding (Fig. S1E and F). Comparison of total TBK1 and phospho-TBK1 (S172) in the starved and fed state revealed that amino acid re-feeding increased both total and phospho-TBK1 on purified lysosomes (Fig. 2 G-I). However, phospho-TBK1 levels in whole cell lysates did not increase noticeably in response to amino acids. This may reflect the presence of only a fraction of the total TBK1 pool at lysosomes. With respect to Rab7, its total abundance at lysosomes was not regulated by either amino acids or TBK1 KO but phospho-Rab7 (S72) levels in both the total lysate and at lysosomes increased in response to amino acid re-feeding and this effect was abolished in cells lacking TBK1 (Fig. 2 G, J and K).

We next took advantage of the robust TBK1-GFP signal to further characterize regulation of lysosomal TBK1 by changes in amino acid availability. Both total and phospho-(S172)-TBK1-GFP increased at lysosomes when starved cells were re-fed with amino acids and this was paralleled by increases in Rab7-S72 phosphorylation (Figure 3A-D). The increase in TBK1-GFP lysosome abundance in response to amino acids was also observed by confocal immunofluorescence microscopy (Fig. 3E). This discovery that TBK1 is dynamically recruited to and signals from lysosomes in response to amino acid abundance, led us to perform additional experiments to validate functionality of TBK1-GFP and to determine the extent to which lysosomes serve as platforms for TBK1 signaling in other contexts. STING is well known to activate TBK1 at the Golgi and to be subsequently degraded within lysosomes (Balka et al., 2023; Gonugunta et al., 2017; Kuchitsu et al., 2023). Consistent with expectations for a functional TBK1 protein, TBK1 KO cells rescued with TBK1-GFP showed strong TBK1 auto-phosphorylation on S172 in response to treatment with cGAMP (STING agonist; Fig. 3F and Fig. S2A). This was also accompanied by an increase in Rab7-S72 and STING-S366 phosphorylation (Fig. 3F and Fig. S2A). Given the predominant late endosome/lysosome localization of Rab7, this implied that TBK1 should also be recruited to lysosomes in response to STING signaling. Indeed, immunoblotting of purified lysosomes from cells treated +/- cGAMP revealed increased lysosome localization of both total and phospho-TBK1 in response to cGAMP treatment (Fig. 3F). Phosphorylated STING also became more abundant at lysosomes in the cGAMP treated cells (Fig. 3F). The increase in both total and phospho-S172-TBK1 at lysosomes following STING activation with cGAMP was also observed by immunofluorescence (Fig. 3G). As a control to further validate the specificity of lysosomal TBK1-GFP detection by immunofluorescence, we observed that both total and phospho-TBK1-GFP showed increased recruitment to lysosomes following lysosome damage induced by LLOME (Fig. 3G)(Eapen et al., 2021; Thiele and Lipsky, 1990). Collectively, these results show that TBK1 undergoes lysosome recruitment and activation in response to multiple activating signals that converge on lysosomes. However, as amino acids still promoted TBK1- dependent Rab7 phosphorylation in STING KO cells, the amino acid dependent pathway for TBK1 activation is independent of STING (Fig. S2B). TBK1 also contributed to Rab7 phosphorylation in response to lysosome damage downstream of LLOME treatment (Fig. S2C).

**Figure 3:**
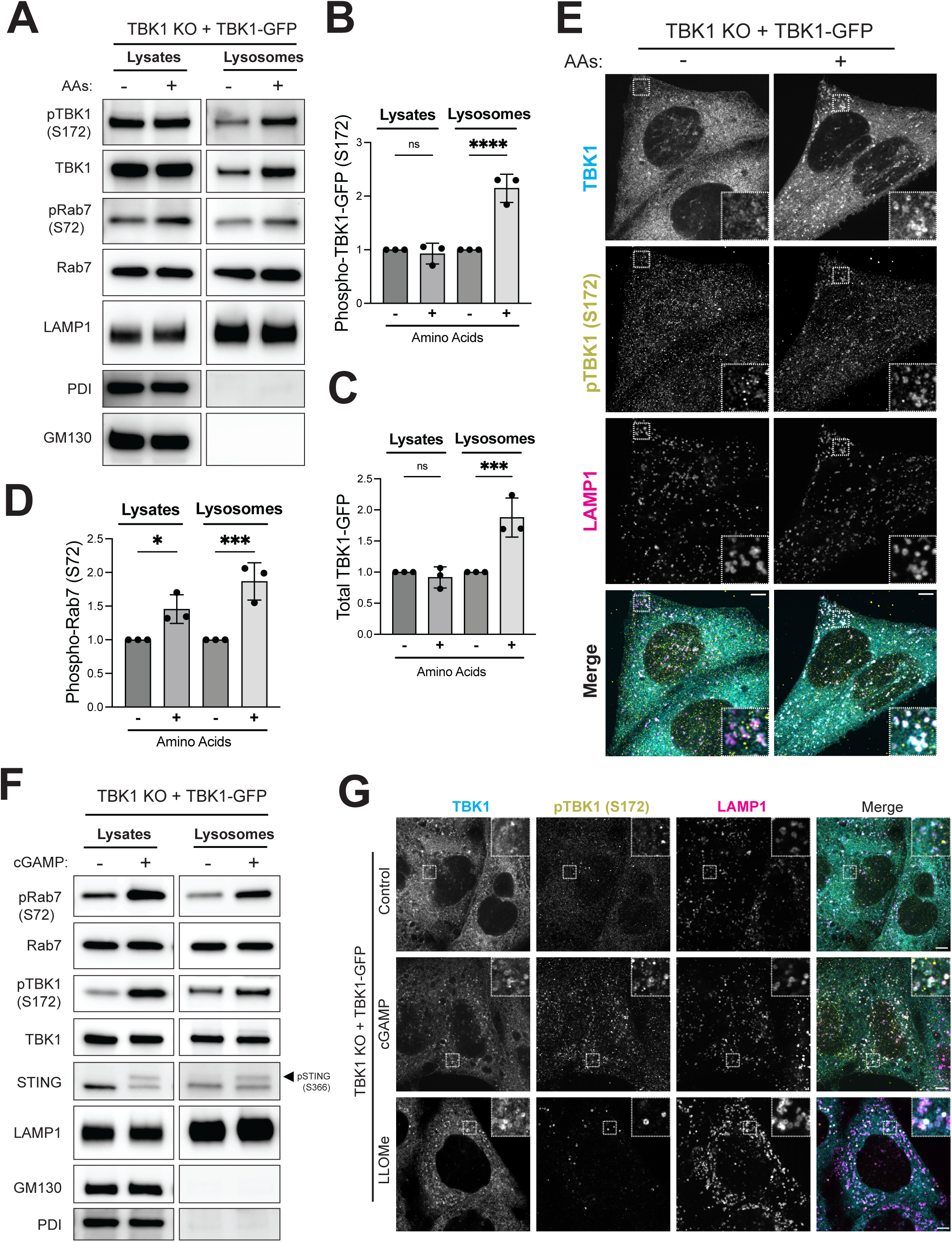
TBK1 is activated and recruited to lysosomes when amino acids are abundant. (A) Immunoblot analysis of the indicated proteins in cell lysates (Lysates) and magnetically isolated lysosomes (Lysosomes) of TBK1 KO stably expressing TBK1-GFP HeLa cells starved for 60 min (-) and then refed with amino acids (3X MEM) for 30’ (+). (B-D) Quantification of lysosomal fold increase by amino acids of phosphorylated TBK1-GFP (pTBK1-GFP) and total TBK1-GFP normalized to total Rab7 and of whole cell lysates and lysosomal fold increase by amino acids of pRab7 (S72) normalized to total Rab7. Data points from three independent experiments (starved cells are considered 1) and statistical significance was determined by ordinary one-way ANOVA with Šidák *post hoc* test (mean ± SD; ****, *p*<0.0001; ***, *p*<0.001; *, *p*<0.05; ns, not significant) (E) Immunofluorescence microscopy analysis of TBK1, pTBK1 (S172) and LAMP1 in TBK1 KO + TBK1-GFP HeLa cells starved for 60 min (-) and re-fed with amino acids (3X) for 30’ (+). (F) Immunoblot analysis of the indicated proteins in TBK1 KO + TBK1-GFP HeLa cells untreated (-) or treated with cGAMP (70 µM) for 120 min (+), shown in cell lysates (Lysates) and magnetically isolated lysosomes (Lysosomes). (G) Immunofluorescence microscopy analysis of TBK1, pTBK1 (S172) and LAMP1 in Control [0.1 % (v/v) DMSO, 30’], cGAMP (70 µM, 2h) or LLOMe (1 mM, 30’) of TBK1 KO + TBK1-GFP HeLa cells. Insets are magnified 3X. Scale bar: 5 µm.

### TBK1-dependent Rab7 phosphorylation is important for efficient mTORC1 activation

Rab7 negatively regulates mTORC1 kinase activity (Kvainickas et al., 2019). However, a role for TBK1-mediated Rab7-S72 phosphorylation in this pathway was not previously reported. In support of a model wherein TBK1-mediated phosphorylation of Rab7 relieves Rab7-dependent inhibition of mTORC1, expression of dominant negative Rab7 (T22N) rescued the impaired mTORC1 activation in TBK1 KO cells (Fig. 4A). Likewise, Rab7 KO on its own increased mTORC1 activation and the double KO of Rab7 + TBK1 rescued the mTORC1 activation defect arising from loss of TBK1 (Fig. 4B and C). The observation that mCherry-tagged wild type Rab7 (but not the nucleotide binding defective T22N mutant) rescued mTORC1 over-activation in Rab7 KO cells, ensured that the relationship between Rab7 and mTORC1 was not an off-target effect of the genome editing or an effect of the clonal selection that was required to generate the KO line (Fig. 4D and S3A).

**Figure 4:**
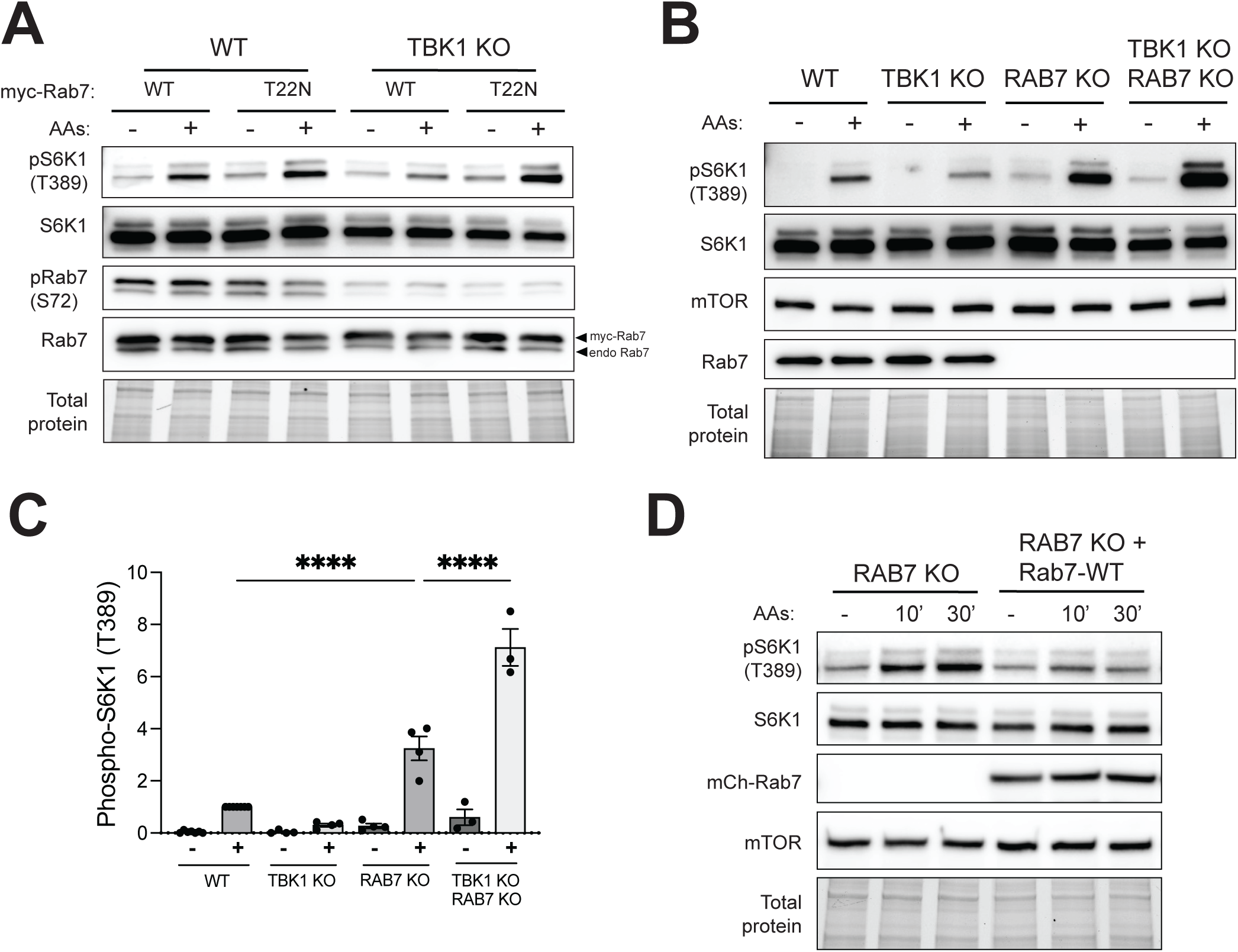
TBK1-Rab7 axis controls mTORC1 signaling. (A) Immunoblotting of the indicated proteins of WT and TBK1 KO HeLa cells transiently expressing myc-tagged Rab7 or Rab7-T22N starved for 60 min (-) and re-fed with amino acids for 60’ (+) (B) Immunoblotting of the indicated proteins of WT, TBK1 KO, RAB7 KO and TBK1 KO RAB7 KO HeLa cells starved for 60 min (-) and re-fed with amino acids for 60 min (+). (C) Quantification of pS6K1 (T389) normalized to total S6K1 shown in B. Plotted data represents at least three independent experiments (WT re-fed with amino acids is considered 1) and statistical significance determined by ordinary one-way ANOVA with Šidák method (mean ± SD; ****, *p*<0.0001). (D) Immunoblotting of the indicated proteins of RAB7 KO and RAB7 KO stably expressing mCherry-tagged wild-type Rab7 (RAB7-WT) of starved cells for 60 min (-) and re-fed with amino acids for 10 min and 30 min (10’ and 30’). TGX stain-free method was used to visualize total protein.

### Rab7-S72 phosphorylation is important for both mTORC1 activation and TBK1 lysosome recruitment

Comparison of Rab7 KO cells that were rescued with equal amounts of wildtype mCherry-Rab7 versus mCherry-Rab7-S72A revealed that this mutant Rab7 that could not be phosphorylated by TBK1 more strongly suppressed mTORC1 activation (Fig. 5A and B). This supports a model where phosphorylation of Rab7 by TBK1 relieves Rab7-dependent suppression of mTORC1 activity. Interestingly, cells expressing the Rab7-S72A mutant had significantly less TBK1 at lysosomes (Fig. 5C and D). The concept that phosphorylation on S72 inhibits major functions of Rab7 is also supported by analysis of STING down-regulation following its activation by cGAMP. This was impaired in Rab7 KO cells, rescued by re-expression of wild type Rab7 and the Rab7-S72A mutant caused even stronger agonist-induced STING degradation (Fig. S3B). These effects of Rab7 phosphorylation on STING degradation match expectations raised by a previous study and thus provide validation for the tools that we have applied to amino acid-dependent regulation of TBK1 at lysosomes (Ritter et al., 2020).

**Figure 5:**
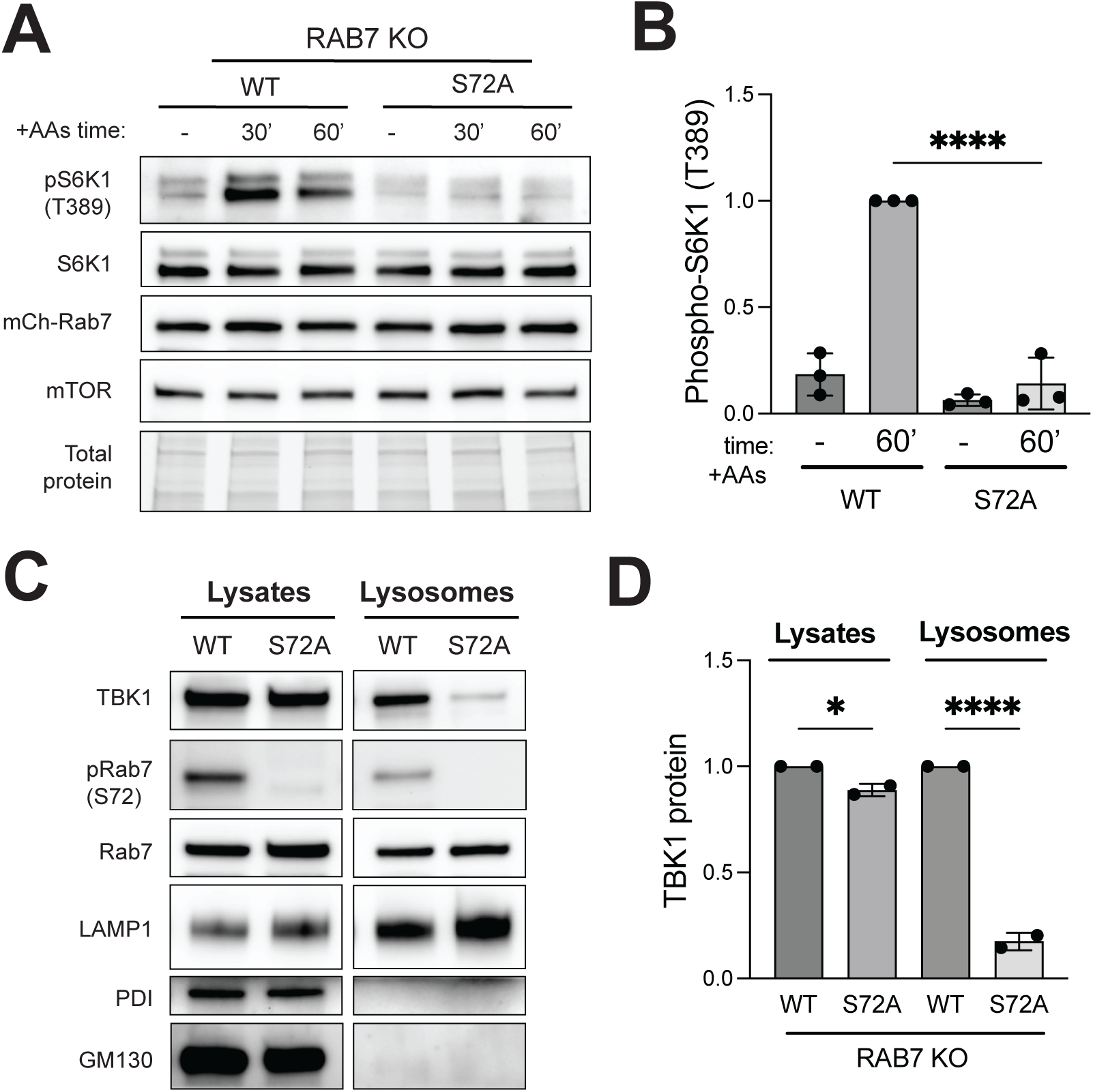
Rab7-S72 phosphorylation is required for efficient mTORC1 activation and TBK1 lysosome localization. (A) Immunoblotting of the indicated proteins of RAB7 KO stably expressing mCherry-tagged wild-type Rab7 (RAB7-WT) or mutant (RAB7-S72A) HeLa cells starved for 60 min (-) and re-fed with amino acids for 30 min and 60 min (30’ and 60’). (B) Quantification of pS6K1 (T389) normalized to total S6K1 in starved cells (-) versus re-fed with amino acids for 60 min (60’). Data points from three independent experiments (re-fed Rab7-WT for 60 min is considered 1) and statistics determined by ordinary one-way ANOVA with Šidák *post hoc* (mean ± SD; ****, *p*<0.0001). TGX stain-free signal was used as a loading control (Total protein). (C) Immunoblot analysis of the indicated proteins in cell lysates (Lysates) and magnetically isolated lysosomes (Lysosomes) of Rab7-WT and Rab7-S72A HeLa cells under basal conditions. (D) TBK1 protein levels were quantified and normalized to Rab7 in the Lysates and Lysosomes, Rab7-WT is considered 1. Statistical analysis was based on ordinary one-way ANOVA with Šidák *post hoc* (mean ± SD; ****, *p*<0.0001; *, *p<*0.05).

**Figure 6:**
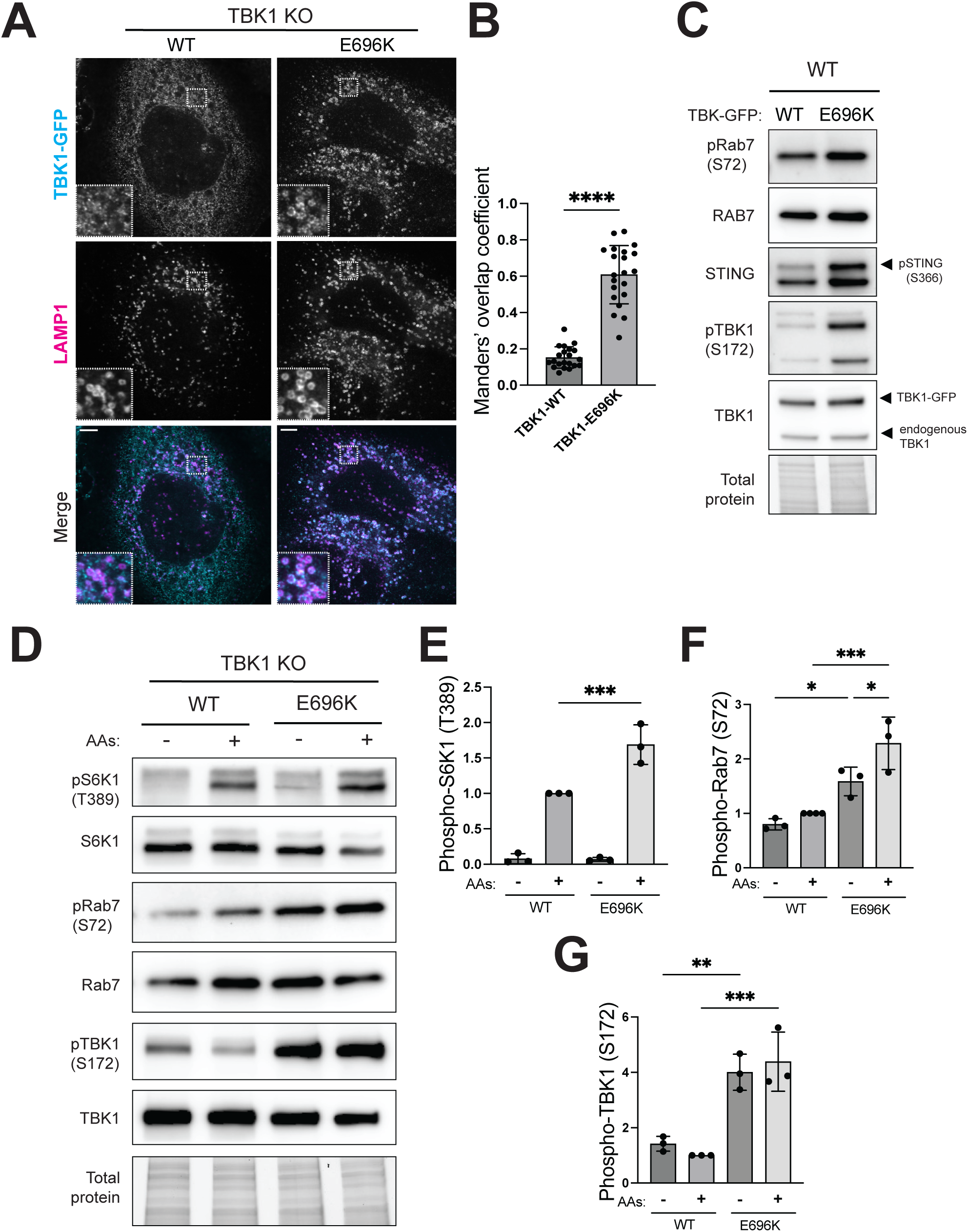
ALS-associated TBK1-E696K mutant accumulates at the lysosomes and over-activates mTORC1. (A) Immunofluorescence microscopy analysis of GFP-tagged wildtype (WT) and mutant (E696K) versions of TBK1 stably expressed in TBK1 KO HeLa cells under basal conditions using GFP and LAMP1 antibodies. TBK1-LAMP1 colocalization analysis based on Manders’ overlap coefficient (M1) and statistical significance determined by unpaired t test (n=3; mean ± SD; ****, two-tailed *p*<0.0001). Each dot represents 1-2 cells (total 30-40 cells). Scale bar: 5 µm. (B) Immunoblot analysis of the indicated proteins in whole cell lysates of TBK1-WT and TBK1-E696K transiently expressed in WT HeLa cells under basal conditions. (C) Immunoblotting of the indicated proteins TBK1-WT and TBK1-E696K stably expressed in TBK1 KO HeLa cells starved for 60 min (-) and re-fed with amino acids for 60 min (+). TGX stain-free method was used to visualize total protein. (D-F) Quantification of phosphorylated proteins normalized to total level and expressed as a fold change compared to the TBK1-WT cells under re-fed conditions. Plotted data are representative of three independent experiments. Statistical significance based on ordinary one-way ANOVA with Šidák *post hoc* test (mean ± SD; ***, *p*<0.001; **, *p*<0.01; *, *p<*0.05).

### TBK1 has Rab7-independent effects on lysosomes

Rab7 has multiple roles in the endolysosomal system and changes in lysosome size occur in response to Rab7 perturbations (Bucci et al., 2000; Kummel et al., 2022). We observed that Rab7 KO and TBK1 KO cells both exhibited a similar increase in late endosome/lysosome size (Fig. 1G, S4A-C). This effect was increased further in Rab7+TBK1 double KO cells (Fig. S4A and B). The lysosome size alterations in TBK1 KO cells were on target as they were rescued by re-introducing TBK1-GFP (Figure S4C). Meanwhile, both wildtype and S72A mutant mCherry-tagged Rab7 proteins localized to LAMP1-positive late endosomes/lysosomes and reversed the increase in lysosome size that occurs in Rab7 KO cells (Fig. S4D and E). However, this effect was more variable with the S72A mutant. The additive effect of the TBK1 and Rab7 KOs and the partial rescue with the Rab7-S72A mutant results indicate that TBK1 likely has additional effects on lysosomes via additional substrates beyond Rab7.

### TBK1-E696K ALS mutant exhibits constitutive lysosome localization, elevated Rab7 phosphorylation and increased mTORC1 activity

TBK1 loss-of-function and missense mutations cause ALS and FTD (Cirulli et al., 2015; Freischmidt et al., 2015; Pottier et al., 2019). It was previously shown that the ALS-associated TBK1-E696K mutant leads to loss-of-function phenotypes related to mitophagy due to reduced interaction with optineurin (Harding et al., 2021; Li et al., 2016; Moore and Holzbaur, 2016). However, it is still unknown if this is the only TBK1-dependent process affected by this mutation. To define the impact of the TBK1-E696K mutation on our newly discovered lysosomal functions of TBK1, we expressed GFP-tagged wild-type or E696K TBK1 (TBK1-WT and TBK1-E696K) in TBK1 KO HeLa cells. Confocal immunofluorescence experiments revealed that stably expressed TBK1-E696K was much more abundant at lysosomes than wild type TBK1 (Fig. 7A and B). This was accompanied by more Rab7-S72 phosphorylation (Fig. 7C). This increased activity for the E696K mutant was not restricted to just Rab7 as there were also increases in TBK1 autophosphorylation and STING phosphorylation in the TBK1-E696K cells under basal growth conditions (Fig. 7C). Increased Rab7 phosphorylation by TBK1-E696K also occurred in the context of amino acid stimulation and was accompanied by more mTORC1 activity (Fig. 7 D-G). Taken together these results indicate that TBK1-E696K has gain-of-function properties as it is hyperactive with respect to lysosome localization, auto-phosphorylation, Rab7 phosphorylation, mTORC1 activation and STING phosphorylation.

## Discussion

TBK1 is well characterized for phosphorylating multiple physiologically important substrates related to innate immunity and organelle quality control (Fitzgerald et al., 2003; Harding et al., 2021; Moore and Holzbaur, 2016; Pied et al., 2022; Weidberg and Elazar, 2011). However, two unbiased phospho-proteomics studies identified Rab7 as a major target for TBK1 (Heo et al., 2018; Ritter et al., 2020). Given that Rab7 localizes to late endosomes and lysosomes this suggested an important but previously unappreciated endolysosomal role (or roles) for TBK1. This furthermore raised questions about both what regulates TBK1 at lysosomes and the physiological consequences of Rab7 phosphorylation. Our results define a nutrient regulated pathway wherein amino acid availability promotes TBK1 activity at lysosomes. By phosphorylating Rab7, this lysosomal TBK1 relieves inhibition of mTORC1 activation by Rab7. The TBK1-E696K mutant that causes ALS-FTD constitutively localizes to lysosomes, phosphorylates Rab7 and enhances mTORC1 amino acid responsiveness. Collectively, these results define a novel pathway linking amino acid availability to the TBK1 and Rab7-dependent regulation of mTORC1 signaling.

Amino acid availability is a major regulator of mTORC1 activation (Lama-Sherpa et al., 2023; Lawrence and Zoncu, 2019; Liu and Sabatini, 2020). This is well appreciated to occur through signals that are sensed and integrated to control the nucleotide loading state of the lysosome-localized Rag GTPase heterodimers that bind mTORC1 and thus control its abundance at lysosomes. Our observations that amino acids also act through TBK1 to control mTORC1 signaling thus represents a novel nutrient input into regulation of the mTORC1 signaling pathway. Placing TBK1 in this nutrient-dependent pathway for mTORC1 activation may be of relevance for the anabolic effects of TBK1 in the context of diabetes and obesity (Bodur et al., 2022; Cruz et al., 2018; Oral et al., 2017; Reilly et al., 2013). Identification of specific amino acids, sensors and signal transduction machinery operating upstream of TBK1 is now a priority. Our elucidation of Rab7 as a TBK1 substrate that controls mTORC1 activation joins previous studies that linked TBK1 to mTORC1 activity via phosphorylation of Raptor and mTOR itself (Antonia et al., 2019; Bodur et al., 2018; Ye et al., 2023). The relative contributions of each of these TBK1 substrates to regulation of mTORC1 activity across physiological contexts remains to be determined.

Rab7 is well established as a regulator of multiple aspects of endolysosomal function including membrane traffic, endosome maturation and lysosome subcellular positioning (Kummel et al., 2023). Rab7 was also previously shown to be a negative regulator of mTORC1 and this effect can be relieved by TBC1D15, a GTPase activating protein (GAP) for Rab7 (Kvainickas et al., 2019). Our results indicate that Rab7-S72 phosphorylation represents an alternative route to relieving the ability of Rab7 to suppress mTORC1.

Leucine rich repeat kinase 2 (LRRK2) phosphorylates multiple members of the Rab family (but not Rab7) on their switch 2 region at a site that corresponds to S72 in Rab7 (Pfeffer, 2022; Steger et al., 2017; Steger et al., 2016). Phosphorylation on this site modulates the regulatory proteins and effectors that these Rabs bind to (Pfeffer, 2022; Steger et al., 2017). This includes a feed forward relationship between LRRK2 and its ability to both phosphorylate Rab8A and Rab10 and to be activated by recruitment to membranes via interactions with phospho-Rab8A and Rab10 (Vides et al., 2022). Our observations that there was significantly less TBK1 at lysosomes in cells that expressed the Rab7-S72A mutant (Fig. 5C and D) suggests the possibility of a similar feed forward role for Rab7 phosphorylation in driving further recruitment of TBK1 to lysosomes.

Although Rab7-S72 phosphorylation and the functional consequences downstream of it are only partially understood, this post-translational modification was previously linked to regulation of macroautophagy, mitophagy, STING down-regulation and EGFR down-regulation (Hanafusa et al., 2019; Heo et al., 2018; Ritter et al., 2020; Tudorica et al., 2023). Consistent with the multiple roles for Rab7 in the endolysosomal pathway, Rab7 phosphorylation is likely to have many parallel effects. Although little is known about Rab7 effectors that differentially respond to phosphorylation on S72, it was very recently proposed that phosphorylation of Rab7 differentially controls its interactions with Rubicon and Pacer, two structurally related effectors with distinct roles in autophagy (Tudorica et al., 2023). This provides a proof of principle for the concept that Rab7 phosphorylation selectively controls effector binding and opens questions about the existence of additional phospho-selective Rab7 effectors that control processes downstream of TBK1 at lysosomes.

The enhanced lysosome localization of the TBK1-E696K mutant provides a valuable tool for investigating the downstream consequences of increasing TBK1 abundance and activity at lysosomes. The constitutive lysosome localization and signaling by the TBK1-E696K mutant also raises the possibility that excessive TBK1 activity at lysosomes contributes to ALS and FTD pathogenesis. To address disease relevance, it would be informative to analyze Rab7 phosphorylation and mTORC1 signaling in samples from patients that carry this mutation. Identifying additional TBK1 mutations that also exhibit increased lysosome localization and Rab7 phosphorylation would also strengthen confidence in the potential disease relevance of the increased TBK1-E696K at lysosomes. However, given that more than 100 TBK1 mutations have so far been discovered in ALS and FTD, it is beyond the scope of this study to systematically test them all (Gurfinkel et al., 2022).

In addition to TBK1, leucine rich repeat kinase 1 (LRRK1) also phosphorylates Rab7 on serine 72 downstream of epidermal growth factor receptor (EGFR) signaling and during mitophagy (Fujita et al., 2022; Hanafusa et al., 2019). This role for LRRK1 likely explains the Rab7 phosphorylation that persisted in the TBK1 KO cells. It remains to be determined whether the pools of Rab7 phosphorylated by TBK1 versus LRRK1 are functionally interchangeable. Identification of the relative contributions of each of these mTORC1 inputs in physiological contexts represents an important next step in this field.

In conclusion, we defined an amino acid regulated TBK1 and Rab7-dependent pathway that is required for efficient mTORC1 activation at lysosomes. This establishes TBK1 as a lysosomal protein that is recruited and activated under conditions of amino acid abundance. By identifying a novel site and regulatory input to TBK1, these results open new avenues for investigating the amino acid sensors acting upstream of TBK1. The demonstration that Rab7 phosphorylation relieves inhibitory effects of Rab7 on mTORC1 activation also raises questions about the identity of relevant effectors that are sensitive to Rab7 phosphorylation state. Finally, the potential contributions of lysosomal TBK1 to ALS and/or FTD pathogenesis is another new research direction arising from the foundation laid by this study.

## Materials and Methods

### Cell culture, growth conditions and drug treatments

HeLa (kindly provided by Pietro De Camilli, Yale University) and RAW 264.7 (ATCC) cells were maintained in DMEM + 10 % FBS + 1 % penicillin/streptomycin (Thermo Fisher Scientific) at 37°C with 5 % CO2. Details of cell lines are summarized in Supplemental Table 1. Experiments to test the effect of amino acids on cells were performed as per our previous studies (Amick et al., 2016; Amick et al., 2020; Meng and Ferguson, 2018; Petit et al., 2013). In brief, cells were washed 2 X in PBS and incubated in RPMI without amino acids (US Biological) for 1 h and then were shifted to RPMI with MEM Amino Acids (Gibco) at the indicated times. BX-795 (Cayman, dissolved in DMSO) and LLOMe (Cayman, dissolved in DMSO) were added to growing cells at the indicated concentrations and compared with cells treated with an equivalent amount of DMSO alone. Details of growth media and drugs are summarized in Supp. Table 2.

### CRISPR/Cas9 genome editing

HeLa TBK1 KO and RAB7 KO cells were created using px459-based Cas9 plasmids (listed in Supp. Table 3). The gRNA sequences to define each target gene are listed in Supp. Table 4. 1 x 10^5^ cells were plated in a 6-well dish per well and transfection mix, containing OPTiMEM, FuGENE HD and 1 µg gRNA plasmid, was added to the cells. RAW 264.7 STING KO and TBK1 + IKKε KO cells were created using the Synthego CRISPR Gene Knockout (Bentley-DeSousa and Ferguson, 2023). 2.5 x 10^5^ cells were plated into a 6-well dish. The following day, cells were transfected with ribonucleoprotein particles using Lipofectamine CRISPRiMAX (Thermo Fisher Scientific), gene specific sgRNA and recombinant Cas9 (Synthego). After 48 h, the media was changed. After 72 h, single cells were plated into 96-well dishes to obtain monoclonal populations, that were further validated by immunoblotting.

### Transposon-mediated genome editing

Transposon-mediated genome editing with piggyBac transposase was used to generate stable HeLa cell lines. 1 x 10^5^ cells were plated per well in a 6-well dish and transfection mix, containing OPTiMEM, FuGENE HD (Promega) and 1:4 ratio of plasmids (0.75 μg piggyBac transposon vector with gene-of-interest + 0.25 μg piggyBac transposase vector) were added to the cells. After 24-48 h, puromycin (Gibco) was added at 2.5 µg/mL. After 24h of selection, surviving cells were recovered in fresh media without puromycin for 24-48 h and then plated into 96-well dishes at 1 cell per well. Monoclonal cell lines were then confirmed by immunoblotting and fluorescence microscopy. Human TBK1-GFP piggyBac vector was purchased from VectorBuilder. mCh-Rab7A was cloned into a piggyBac plasmid using the NEB HIFI DNA Assembly Kit (NEB) and primers listed in Supp. Table 4. mCh-Rab7A was a gift from Gia Voeltz (Addgene plasmid #61804; http://n2t.net/addgene:61804; RRID:Addgene_61804). Site-directed mutagenesis of TBK1 and Rab7 genes was performed using Q5 High-Fidelity 2X Master Mix (NEB). Plasmids are listed in Table 3 and were confirmed by Sanger Sequencing (Keck DNA Sequencing facility, Yale University).

### Transient transfections

3 x 10^5^ HeLa cells were plated in a 6-well dish per well overnight. Next day, a transfection mix, containing 200 µL OPTiMEM, 6 µL Lipofectamine 2000 (Invitrogen) and 2 µg plasmid, per well, was added to the cells for 24 h.

### Immunoblotting

3 x 10^5^ HeLa or 1 x 10^6^ RAW 264.7 cells were seeded per well in a 6-well dish. The following day, cells were washed 2X with cold PBS and scraped in cold lysis buffer [50 mM Tris-HCl pH 7.4, 150 mM NaCl, 1 % (v/v) Triton X-100, 1 mM EDTA supplemented with protease and phosphatase inhibitors]. Lysates were centrifuged at 20,817 x g (4 °C) for 8 min to pellet and discard insoluble material. Protein concentrations were measured by Bradford method (Coomassie Plus Protein Assay Reagent, ThermoFisher Scientific). The lysate supernatants were mixed 1:1 with Laemmli Buffer [80 mM Tris-HCl pH 6.8, 3.4 M Glycerol, 3 % (w/v) SDS, and Bromophenol Blue supplemented with fresh 6 % (v/v) β-mercaptoethanol] and heated at 95 °C for 3 min.

Protein lysates were electrophoresed in 4-15 % miniPROTEAN TGX Stain-Free pre-cast gels (BioRad) using electrophoresis buffer [25 mM Tris Base, 192 mM Glycine, 0.1 % (w/v) SDS. After electrophoresis, gels were transferred onto 0.22 µm pore nitrocellulose membrane (Thermo Fisher Scientific) at 100 V for 60 min in transfer buffer (25 mM Tris Base, 192 mM Glycine, 20 % (v/v) Methanol). Total protein was visualized using BioRad TGX stain-free technology prior to blocking. Membranes were blocked in 5 % non-fat dry milk in TBS-T [10 mM Tris Base, 150 mM NaCl, 0.1 % (v/v) Tween 20]. Antibodies were added to the membranes in 5 % (w/v) BSA (Sigma-Aldrich) in TBS-T overnight (4 °C) at the indicated concentrations (Supp. Table 5). Membranes were washed 3X for 10 minutes with TBS-T. Secondary antibodies were added in TBS-T or 5% (w/v) non-fat dry milk TBS-T for 1 h at RT. Membranes were washed 3X for 10 min with TBS-T. Membranes were subjected to chemiluminescence by adding Pico ECL or Femto ECL (Thermo Fisher Scientific) and visualized in a BioRad Chemidoc MP imaging station.

### Immunoprecipitation

TBK1-GFP expressed in HeLa TBK1 KO cells was immunoprecipitated to analyze TBK1-STING interaction before and after cGAMP addition. Briefly, cells were washed 2X with cold PBS-1X, scraped in ice-cold lysis buffer mentioned above, and centrifuged at 20,817 x g (4 °C) for 8 min. Protein concentrations were measured by Bradford method and lysates were then incubated with rotation in pre-washed GFP-TRAP beads (Chromotek) for 1 h at 4 °C. Beads were washed 5X with lysis buffer. Proteins were eluted by 0.5X Laemmli Buffer and boiled at 95 °C for 3 min.

### Immunofluorescence microscopy

Immunofluorescence microscopy was performed with a Nikon Ti2-E inverted microscope equipped with Spinning Disk Super Resolution by Optical Pixel Reassignment Microscope (Yokogawa CSU-W1 SoRa, Nikon) and Microlens-enhanced Nipkow Disk with pinholes and 60x SR Plan Apo IR oil immersion objective. Images were acquired at room temperature. Cells were grown on 12 mm coverslips (GmbH & Co KG), washed in PBS and fixed by adding 4 % PFA in 0.1 M sodium phosphate. Cells were fixed at room temperature for 25 min and then washed 3X with PBS. Samples were permeabilized by immersing coverslips in ice-cold methanol for 2-3 s, followed by PBS rinses. Samples were then blocked in PBS-1X with 5% normal donkey serum (Jackson) for 1 h at RT. Subsequent antibody incubations were performed in this buffer. Antibodies used in this study are listed in Table 5.

Images were processed with ImageJ/FIJI (Schindelin et al., 2012). For analysis of protein colocalization, Manders’ coefficient was determined with ImageJ/Fiji through JACoP plugin (Bolte and Cordelieres, 2006). For lysosome size measurements, lysosome perimeter was determined with ImageJ/Fiji through StarDist plugin (Schmidt et al., 2018). Thresholds were the same for all images analyzed in a given experiment.

### Lysosome isolation with SPIONs

The magnetic isolation of lysosomes was performed by endocytic loading of cells with superparamagnetic iron oxide nanoparticles (SPIONs) that were prepared as previously described (Amick et al., 2018; Rodriguez-Paris et al., 1993; Tharkeshwar et al., 2020). In short, 2.5 x 10^7^ HeLa cells were plated in a 15 cm dish overnight. 10% SPION-containing media was added to the cells for 24 h. Cells were then trypsinized and plated in 15-cm dishes at 2.5 x 10^7^ for 18 h in fresh media that lacked SPIONs.

### Statistical analysis

Statistical analysis was carried out with GraphPad Prism version 10.1.0, with specific details about the statistical tests conducted, the number of independent experiments, and *p* values provided in the corresponding figure legends.

## Acknowledgements

This research was supported by grants from the National Institutes for Health (GM105718) and Aligning Science Across Parkinson’s disease (ASAP-000580) through the Michael J. Fox Foundation for Parkinson’s Research (MJFF). Agnes Roczniak-Ferguson (Yale) provided valuable feedback and key plasmids. Joe Amick (Yale) generated plasmids for Rab7 knockout. We thank Berrak Ugur (Yale) for managing our ASAP project. The authors do not have any conflicts to declare.

## Author contributions

GT and SMF designed experiments. GT and ABD performed experiments. GT and SMF analyzed data and prepared the manuscript.

## References

Ahmad, L., S.Y. Zhang, J.L. Casanova, and V. Sancho-Shimizu. 2016. Human TBK1: A Gatekeeper of Neuroinflammation. Trends Mol Med. 22:511–527.

Amick, J., A. Roczniak-Ferguson, and S.M. Ferguson. 2016. C9orf72 binds SMCR8, localizes to lysosomes, and regulates mTORC1 signaling. Mol Biol Cell. 27:3040–3051.

Amick, J., A.K. Tharkeshwar, C. Amaya, and S.M. Ferguson. 2018. WDR41 supports lysosomal response to changes in amino acid availability. Mol Biol Cell. 29:2213–2227.

Amick, J., A.K. Tharkeshwar, G. Talaia, and S.M. Ferguson. 2020. PQLC2 recruits the C9orf72 complex to lysosomes in response to cationic amino acid starvation. J Cell Biol. 219.

Angarola, B., and S.M. Ferguson. 2020. Coordination of Rheb lysosomal membrane interactions with mTORC1 activation. F1000Res. 9.

Antonia, R.J., J. Castillo, L.E. Herring, D.S. Serafin, P. Liu, L.M. Graves, A.S. Baldwin, and R.S. Hagan. 2019. TBK1 Limits mTORC1 by Promoting Phosphorylation of Raptor Ser877. Sci Rep. 9:13470.

Bain, J., L. Plater, M. Elliott, N. Shpiro, C.J. Hastie, H. McLauchlan, I. Klevernic, J.S. Arthur, D.R. Alessi, and P. Cohen. 2007. The selectivity of protein kinase inhibitors: a further update. Biochem J. 408:297–315.

Balka, K.R., C. Louis, T.L. Saunders, A.M. Smith, D.J. Calleja, D.B. D’Silva, F. Moghaddas, M. Tailler, K.E. Lawlor, Y. Zhan, C.J. Burns, I.P. Wicks, J.J. Miner, B.T. Kile, S.L. Masters, and D. De Nardo. 2020. TBK1 and IKKepsilon Act Redundantly to Mediate STING-Induced NF-kappaB Responses in Myeloid Cells. Cell Rep. 31:107492.

Balka, K.R., R. Venkatraman, T.L. Saunders, A. Shoppee, E.S. Pang, Z. Magill, J. Homman-Ludiye, C. Huang, R.M. Lane, H.M. York, P. Tan, R.B. Schittenhelm, S. Arumugam, B.T. Kile, M. O’Keeffe, and D. De Nardo. 2023. Termination of STING responses is mediated via ESCRT-dependent degradation. EMBO J. 42:e112712.

Ballabio, A., and J.S. Bonifacino. 2019. Lysosomes as dynamic regulators of cell and organismal homeostasis. Nat Rev Mol Cell Biol.

Bentley-DeSousa, A., and S.M. Ferguson. 2023. A STING-CASM-GABARAP Pathway Activates LRRK2 at Lysosomes. bioRxiv.

Bodur, C., D. Kazyken, K. Huang, B. Ekim Ustunel, K.A. Siroky, A.S. Tooley, I.E. Gonzalez, D.H. Foley, H.A. Acosta-Jaquez, T.M. Barnes, G.K. Steinl, K.W. Cho, C.N. Lumeng, S.M. Riddle, M.G. Myers, Jr., and D.C. Fingar. 2018. The IKK-related kinase TBK1 activates mTORC1 directly in response to growth factors and innate immune agonists. EMBO J. 37:19–38.

Bodur, C., D. Kazyken, K. Huang, A.S. Tooley, K.W. Cho, T.M. Barnes, C.N. Lumeng, M.G. Myers, and D.C. Fingar. 2022. TBK1-mTOR Signaling Attenuates Obesity-Linked Hyperglycemia and Insulin Resistance. Diabetes. 71:2297–2312.

Bohannon, K.P., and P.I. Hanson. 2020. ESCRT puts its thumb on the nanoscale: Fixing tiny holes in endolysosomes. Curr Opin Cell Biol. 65:122–130.

Bolte, S., and F.P. Cordelieres. 2006. A guided tour into subcellular colocalization analysis in light microscopy. J Microsc. 224:213–232.

Bucci, C., P. Thomsen, P. Nicoziani, J. McCarthy, and B. van Deurs. 2000. Rab7: a key to lysosome biogenesis. Mol Biol Cell. 11:467–480.

Cirulli, E.T., B.N. Lasseigne, S. Petrovski, P.C. Sapp, P.A. Dion, C.S. Leblond, J. Couthouis, Y.F. Lu, Q. Wang, B.J. Krueger, Z. Ren, J. Keebler, Y. Han, S.E. Levy, B.E. Boone, J.R. Wimbish, L.L. Waite, A.L. Jones, J.P. Carulli, A.G. Day-Williams, J.F. Staropoli, W.W. Xin, A. Chesi, A.R. Raphael, D. McKenna-Yasek, J. Cady, J.M. Vianney de Jong, K.P. Kenna, B.N. Smith, S. Topp, J. Miller, A. Gkazi, F.S. Consortium, A. Al-Chalabi, L.H. van den Berg, J. Veldink, V. Silani, N. Ticozzi, C.E. Shaw, R.H. Baloh, S. Appel, E. Simpson, C. Lagier-Tourenne, S.M. Pulst, S. Gibson, J.Q. Trojanowski, L. Elman, L. McCluskey, M. Grossman, N.A. Shneider, W.K. Chung, J.M. Ravits, J.D. Glass, K.B. Sims, V.M. Van Deerlin, T. Maniatis, S.D. Hayes, A. Ordureau, S. Swarup, J. Landers, F. Baas, A.S. Allen, R.S. Bedlack, J.W. Harper, A.D. Gitler, G.A. Rouleau, R. Brown, M.B. Harms, G.M. Cooper, T. Harris, R.M. Myers, and D.B. Goldstein. 2015. Exome sequencing in amyotrophic lateral sclerosis identifies risk genes and pathways. Science. 347:1436–1441.

Cooper, J.M., Y.H. Ou, E.A. McMillan, R.M. Vaden, A. Zaman, B.O. Bodemann, G. Makkar, B.A. Posner, and M.A. White. 2017. TBK1 Provides Context-Selective Support of the Activated AKT/mTOR Pathway in Lung Cancer. Cancer Res. 77:5077–5094.

Cruz, V.H., E.N. Arner, K.W. Wynne, P.E. Scherer, and R.A. Brekken. 2018. Loss of Tbk1 kinase activity protects mice from diet-induced metabolic dysfunction. Mol Metab. 16:139–149.

Eapen, V.V., S. Swarup, M.J. Hoyer, J.A. Paulo, and J.W. Harper. 2021. Quantitative proteomics reveals the selectivity of ubiquitin-binding autophagy receptors in the turnover of damaged lysosomes by lysophagy. Elife. 10.

Ferguson, S.M. 2015. Beyond indigestion: emerging roles for lysosome-based signaling in human disease. Curr Opin Cell Biol. 35:59–68.

Fitzgerald, K.A., S.M. McWhirter, K.L. Faia, D.C. Rowe, E. Latz, D.T. Golenbock, A.J. Coyle, S.M. Liao, and T. Maniatis. 2003. IKKepsilon and TBK1 are essential components of the IRF3 signaling pathway. Nat Immunol. 4:491–496.

Freischmidt, A., T. Wieland, B. Richter, W. Ruf, V. Schaeffer, K. Muller, N. Marroquin, F. Nordin, A. Hubers, P. Weydt, S. Pinto, R. Press, S. Millecamps, N. Molko, E. Bernard, C. Desnuelle, M.H. Soriani, J. Dorst, E. Graf, U. Nordstrom, M.S. Feiler, S. Putz, T.M. Boeckers, T. Meyer, A.S. Winkler, J. Winkelman, M. de Carvalho, D.R. Thal, M. Otto, T. Brannstrom, A.E. Volk, P. Kursula, K.M. Danzer, P. Lichtner, I. Dikic, T. Meitinger, A.C. Ludolph, T.M. Strom, P.M. Andersen, and J.H. Weishaupt. 2015. Haploinsufficiency of TBK1 causes familial ALS and fronto-temporal dementia. Nat Neurosci. 18:631–636.

Fujita, K., S. Kedashiro, T. Yagi, N. Hisamoto, K. Matsumoto, and H. Hanafusa. 2022. The ULK complex-LRRK1 axis regulates Parkin-mediated mitophagy via Rab7 Ser-72 phosphorylation. J Cell Sci. 135.

Gijselinck, I., S. Van Mossevelde, J. van der Zee, A. Sieben, S. Philtjens, B. Heeman, S. Engelborghs, M. Vandenbulcke, G. De Baets, V. Baumer, I. Cuijt, M. Van den Broeck, K. Peeters, M. Mattheijssens, F. Rousseau, R. Vandenberghe, P. De Jonghe, P. Cras, P.P. De Deyn, J.J. Martin, M. Cruts, C. Van Broeckhoven, and B. Consortium. 2015. Loss of TBK1 is a frequent cause of frontotemporal dementia in a Belgian cohort. Neurology. 85:2116–2125.

Gonugunta, V.K., T. Sakai, V. Pokatayev, K. Yang, J. Wu, N. Dobbs, and N. Yan. 2017. Trafficking-Mediated STING Degradation Requires Sorting to Acidified Endolysosomes and Can Be Targeted to Enhance Anti-tumor Response. Cell Rep. 21:3234–3242.

Goul, C., R. Peruzzo, and R. Zoncu. 2023. The molecular basis of nutrient sensing and signalling by mTORC1 in metabolism regulation and disease. Nat Rev Mol Cell Biol. 24:857–875.

Gurfinkel, Y., N. Polain, K. Sonar, P. Nice, R.L. Mancera, and S.L. Rea. 2022. Functional and structural consequences of TBK1 missense variants in frontotemporal lobar degeneration and amyotrophic lateral sclerosis. Neurobiol Dis. 174:105859.

Hanafusa, H., T. Yagi, H. Ikeda, N. Hisamoto, T. Nishioka, K. Kaibuchi, K. Shirakabe, and K. Matsumoto. 2019. LRRK1 phosphorylation of Rab7 at S72 links trafficking of EGFR-containing endosomes to its effector RILP. J Cell Sci. 132.

Harding, O., C.S. Evans, J. Ye, J. Cheung, T. Maniatis, and E.L.F. Holzbaur. 2021. ALS- and FTD-associated missense mutations in TBK1 differentially disrupt mitophagy. Proc Natl Acad Sci U S A. 118.

Hasan, M., V.K. Gonugunta, N. Dobbs, A. Ali, G. Palchik, M.A. Calvaruso, R.J. DeBerardinis, and N. Yan. 2017. Chronic innate immune activation of TBK1 suppresses mTORC1 activity and dysregulates cellular metabolism. Proc Natl Acad Sci U S A. 114:746–751.

Heo, J.M., A. Ordureau, S. Swarup, J.A. Paulo, K. Shen, D.M. Sabatini, and J.W. Harper. 2018. RAB7A phosphorylation by TBK1 promotes mitophagy via the PINK-PARKIN pathway. Sci Adv. 4:eaav0443.

Hirsch-Reinshagen, V., O.A. Alfaify, G.R. Hsiung, C. Pottier, M. Baker, R.B. Perkerson, 3rd, R. Rademakers, H. Briemberg, D.J. Foti, and I.R. Mackenzie. 2019. Clinicopathologic correlations in a family with a TBK1 mutation presenting as primary progressive aphasia and primary lateral sclerosis. Amyotroph Lateral Scler Frontotemporal Degener. 20:568–575.

Hung, Y.H., L.M. Chen, J.Y. Yang, and W.Y. Yang. 2013. Spatiotemporally controlled induction of autophagy-mediated lysosome turnover. Nat Commun. 4:2111.

Kuchitsu, Y., K. Mukai, R. Uematsu, Y. Takaada, A. Shinojima, R. Shindo, T. Shoji, S. Hamano, E. Ogawa, R. Sato, K. Miyake, A. Kato, Y. Kawaguchi, M. Nishitani-Isa, K. Izawa, R. Nishikomori, T. Yasumi, T. Suzuki, N. Dohmae, T. Uemura, G.N. Barber, H. Arai, S. Waguri, and T. Taguchi. 2023. STING signalling is terminated through ESCRT-dependent microautophagy of vesicles originating from recycling endosomes. Nat Cell Biol. 25:453–466.

Kummel, D., E. Herrmann, L. Langemeyer, and C. Ungermann. 2022. Molecular insights into endolysosomal microcompartment formation and maintenance. Biol Chem.

Kummel, D., E. Herrmann, L. Langemeyer, and C. Ungermann. 2023. Molecular insights into endolysosomal microcompartment formation and maintenance. Biol Chem. 404:441–454.

Kvainickas, A., H. Nagele, W. Qi, L. Dokladal, A. Jimenez-Orgaz, L. Stehl, D. Gangurde, Q. Zhao, Z. Hu, J. Dengjel, C. De Virgilio, R. Baumeister, and F. Steinberg. 2019. Retromer and TBC1D5 maintain late endosomal RAB7 domains to enable amino acid-induced mTORC1 signaling. J Cell Biol. 218:3019–3038.

Lama-Sherpa, T.D., M.H. Jeong, and J.L. Jewell. 2023. Regulation of mTORC1 by the Rag GTPases. Biochem Soc Trans. 51:655–664.

Lawrence, R.E., and R. Zoncu. 2019. The lysosome as a cellular centre for signalling, metabolism and quality control. Nat Cell Biol. 21:133–142.

Li, F., X. Xie, Y. Wang, J. Liu, X. Cheng, Y. Guo, Y. Gong, S. Hu, and L. Pan. 2016. Structural insights into the interaction and disease mechanism of neurodegenerative disease-associated optineurin and TBK1 proteins. Nat Commun. 7:12708.

Liu, G.Y., and D.M. Sabatini. 2020. mTOR at the nexus of nutrition, growth, ageing and disease. Nat Rev Mol Cell Biol.

Maejima, I., A. Takahashi, H. Omori, T. Kimura, Y. Takabatake, T. Saitoh, A. Yamamoto, M. Hamasaki, T. Noda, Y. Isaka, and T. Yoshimori. 2013. Autophagy sequesters damaged lysosomes to control lysosomal biogenesis and kidney injury. EMBO J. 32:2336–2347.

Meng, J., and S.M. Ferguson. 2018. GATOR1-dependent recruitment of FLCN-FNIP to lysosomes coordinates Rag GTPase heterodimer nucleotide status in response to amino acids. J Cell Biol. 217:2765–2776.

Menon, S., C.C. Dibble, G. Talbott, G. Hoxhaj, A.J. Valvezan, H. Takahashi, L.C. Cantley, and B.D. Manning. 2014. Spatial control of the TSC complex integrates insulin and nutrient regulation of mTORC1 at the lysosome. Cell. 156:771–785.

Moore, A.S., and E.L. Holzbaur. 2016. Dynamic recruitment and activation of ALS-associated TBK1 with its target optineurin are required for efficient mitophagy. Proc Natl Acad Sci U S A. 113:E3349–3358.

Oral, E.A., S.M. Reilly, A.V. Gomez, R. Meral, L. Butz, N. Ajluni, T.L. Chenevert, E. Korytnaya, A.H. Neidert, R. Hench, D. Rus, J.F. Horowitz, B. Poirier, P. Zhao, K. Lehmann, M. Jain, R. Yu, C. Liddle, M. Ahmadian, M. Downes, R.M. Evans, and A.R. Saltiel. 2017. Inhibition of IKKvarepsilon and TBK1 Improves Glucose Control in a Subset of Patients with Type 2 Diabetes. Cell Metab. 26:157–170 e157.

Petit, C.S., A. Roczniak-Ferguson, and S.M. Ferguson. 2013. Recruitment of folliculin to lysosomes supports the amino acid-dependent activation of Rag GTPases. J Cell Biol. 202:1107–1122.

Pfeffer, S.R. 2022. LRRK2 phosphorylation of Rab GTPases in Parkinson’s disease. FEBS Lett.

Pied, N., C.F. Daussy, Z. Denis, J. Ragues, M. Faure, R. Iggo, M.P. Tschan, B. Roger, F. Rayne, and H. Wodrich. 2022. TBK1 is part of a galectin 8 dependent membrane damage recognition complex and drives autophagy upon Adenovirus endosomal escape. PLoS Pathog. 18:e1010736.

Pomerantz, J.L., and D. Baltimore. 1999. NF-kappaB activation by a signaling complex containing TRAF2, TANK and TBK1, a novel IKK-related kinase. EMBO J. 18:6694–6704.

Pottier, C., Y. Ren, R.B. Perkerson, 3rd, M. Baker, G.D. Jenkins, M. van Blitterswijk, M. DeJesus-Hernandez, J.G.J. van Rooij, M.E. Murray, E. Christopher, S.K. McDonnell, Z. Fogarty, A. Batzler, S. Tian, C.T. Vicente, B. Matchett, A.M. Karydas, G.R. Hsiung, H. Seelaar, M.O. Mol, E.C. Finger, C. Graff, L. Oijerstedt, M. Neumann, P. Heutink, M. Synofzik, C. Wilke, J. Prudlo, P. Rizzu, J. Simon-Sanchez, D. Edbauer, S. Roeber, J. Diehl-Schmid, B.M. Evers, A. King, M.M. Mesulam, S. Weintraub, C. Geula, K.F. Bieniek, L. Petrucelli, G.L. Ahern, E.M. Reiman, B.K. Woodruff, R.J. Caselli, E.D. Huey, M.R. Farlow, J. Grafman, S. Mead, L.T. Grinberg, S. Spina, M. Grossman, D.J. Irwin, E.B. Lee, E. Suh, J. Snowden, D. Mann, N. Ertekin-Taner, R.J. Uitti, Z.K. Wszolek, K.A. Josephs, J.E. Parisi, D.S. Knopman, R.C. Petersen, J.R. Hodges, O. Piguet, E.G. Geier, J.S. Yokoyama, R.A. Rissman, E. Rogaeva, J. Keith, L. Zinman, M.C. Tartaglia, N.J. Cairns, C. Cruchaga, B. Ghetti, J. Kofler, O.L. Lopez, T.G. Beach, T. Arzberger, J. Herms, L.S. Honig, J.P. Vonsattel, G.M. Halliday, J.B. Kwok, C.L. White, 3rd, M. Gearing, J. Glass, S. Rollinson, S. Pickering-Brown, J.D. Rohrer, J.Q. Trojanowski, V. Van Deerlin, E.H. Bigio, C. Troakes, S. Al-Sarraj, Y. Asmann, B.L. Miller, N.R. Graff-Radford, B.F. Boeve, W.W. Seeley, et al. 2019. Genome-wide analyses as part of the international FTLD-TDP whole-genome sequencing consortium reveals novel disease risk factors and increases support for immune dysfunction in FTLD. Acta Neuropathol. 137:879–899.

Reilly, S.M., S.H. Chiang, S.J. Decker, L. Chang, M. Uhm, M.J. Larsen, J.R. Rubin, J. Mowers, N.M. White, I. Hochberg, M. Downes, R.T. Yu, C. Liddle, R.M. Evans, D. Oh, P. Li, J.M. Olefsky, and A.R. Saltiel. 2013. An inhibitor of the protein kinases TBK1 and IKK-varepsilon improves obesity-related metabolic dysfunctions in mice. Nat Med. 19:313–321.

Ritter, J.L., Z. Zhu, T.C. Thai, N.R. Mahadevan, P. Mertins, E.H. Knelson, B.P. Piel, S. Han, J.D. Jaffe, S.A. Carr, D.A. Barbie, and T.U. Barbie. 2020. Phosphorylation of RAB7 by TBK1/IKKepsilon Regulates Innate Immune Signaling in Triple-Negative Breast Cancer. Cancer Res. 80:44–56.

Rodriguez-Paris, J.M., K.V. Nolta, and T.L. Steck. 1993. Characterization of lysosomes isolated from Dictyostelium discoideum by magnetic fractionation. J Biol Chem. 268:9110–9116.

Schindelin, J., I. Arganda-Carreras, E. Frise, V. Kaynig, M. Longair, T. Pietzsch, S. Preibisch, C. Rueden, S. Saalfeld, B. Schmid, J.Y. Tinevez, D.J. White, V. Hartenstein, K. Eliceiri, P. Tomancak, and A. Cardona. 2012. Fiji: an open-source platform for biological-image analysis. *Nat Methods*. 9:676-682.

Schmidt, U., M. Weigert, C. Broaddus, and G. Myers. 2018. Cell Detection with Star-Convex Polygons. Medical Image Computing and Computer Assisted Intervention – MICCAI 2018.

Steger, M., F. Diez, H.S. Dhekne, P. Lis, R.S. Nirujogi, O. Karayel, F. Tonelli, T.N. Martinez, E. Lorentzen, S.R. Pfeffer, D.R. Alessi, and M. Mann. 2017. Systematic proteomic analysis of LRRK2-mediated Rab GTPase phosphorylation establishes a connection to ciliogenesis. Elife. 6.

Steger, M., F. Tonelli, G. Ito, P. Davies, M. Trost, M. Vetter, S. Wachter, E. Lorentzen, G. Duddy, S. Wilson, M.A. Baptista, B.K. Fiske, M.J. Fell, J.A. Morrow, A.D. Reith, D.R. Alessi, and M. Mann. 2016. Phosphoproteomics reveals that Parkinson’s disease kinase LRRK2 regulates a subset of Rab GTPases. Elife. 5.

Tanaka, Y., and Z.J. Chen. 2012. STING specifies IRF3 phosphorylation by TBK1 in the cytosolic DNA signaling pathway. Sci Signal. 5:ra20.

Tharkeshwar, A.K., D. Demedts, and W. Annaert. 2020. Superparamagnetic Nanoparticles for Lysosome Isolation to Identify Spatial Alterations in Lysosomal Protein and Lipid Composition. STAR Protoc. 1:100122.

Thiele, D.L., and P.E. Lipsky. 1990. Mechanism of L-leucyl-L-leucine methyl ester-mediated killing of cytotoxic lymphocytes: dependence on a lysosomal thiol protease, dipeptidyl peptidase I, that is enriched in these cells. Proc Natl Acad Sci U S A. 87:83–87.

Tudorica, D.A., B. Basak, A.P. Cordova, G. Khuu, K. Rose, M. Lazarou, E.L. Holzbaur, and J.H. Hurley. 2023. A RAB7A Phosphoswitch Coordinates Rubicon Homology Protein Regulation of PINK1/Parkin-Dependent Mitophagy. BioRXiv.

Van Mossevelde, S., J. van der Zee, I. Gijselinck, S. Engelborghs, A. Sieben, T. Van Langenhove, J. De Bleecker, J. Baets, M. Vandenbulcke, K. Van Laere, S. Ceyssens, M. Van den Broeck, K. Peeters, M. Mattheijssens, P. Cras, R. Vandenberghe, P. De Jonghe, J.J. Martin, P.P. De Deyn, M. Cruts, C. Van Broeckhoven, and c. Belgian Neurology. 2016. Clinical features of TBK1 carriers compared with C9orf72, GRN and non-mutation carriers in a Belgian cohort. Brain. 139:452–467.

Vargas, J.N.S., M. Hamasaki, T. Kawabata, R.J. Youle, and T. Yoshimori. 2023. The mechanisms and roles of selective autophagy in mammals. Nat Rev Mol Cell Biol. 24:167–185.

Vides, E.G., A. Adhikari, C.Y. Chiang, P. Lis, E. Purlyte, C. Limouse, J.L. Shumate, E. Spinola-Lasso, H.S. Dhekne, D.R. Alessi, and S.R. Pfeffer. 2022. A feed-forward pathway drives LRRK2 kinase membrane recruitment and activation. Elife. 11.

Weidberg, H., and Z. Elazar. 2011. TBK1 mediates crosstalk between the innate immune response and autophagy. Sci Signal. 4:pe39.

Yang, H., and J.X. Tan. 2023. Lysosomal quality control: molecular mechanisms and therapeutic implications. Trends Cell Biol. 33:749–764.

Ye, M., Y. Hu, B. Zhao, Q. Mou, Y. Ni, J. Luo, L. Li, H. Zhang, and Y. Zhao. 2023. TBK1 Knockdown Alleviates Axonal Transport Deficits in Retinal Ganglion Cells Via mTORC1 Activation in a Retinal Damage Mouse Model. Invest Ophthalmol Vis Sci. 64:1.

