## Supplemental Data for "Lysosomal TBK1 Responds to Amino Acid Availability to Relieve Rab7-Dependent mTORC1 Inhibition"

### Supplemental Figures

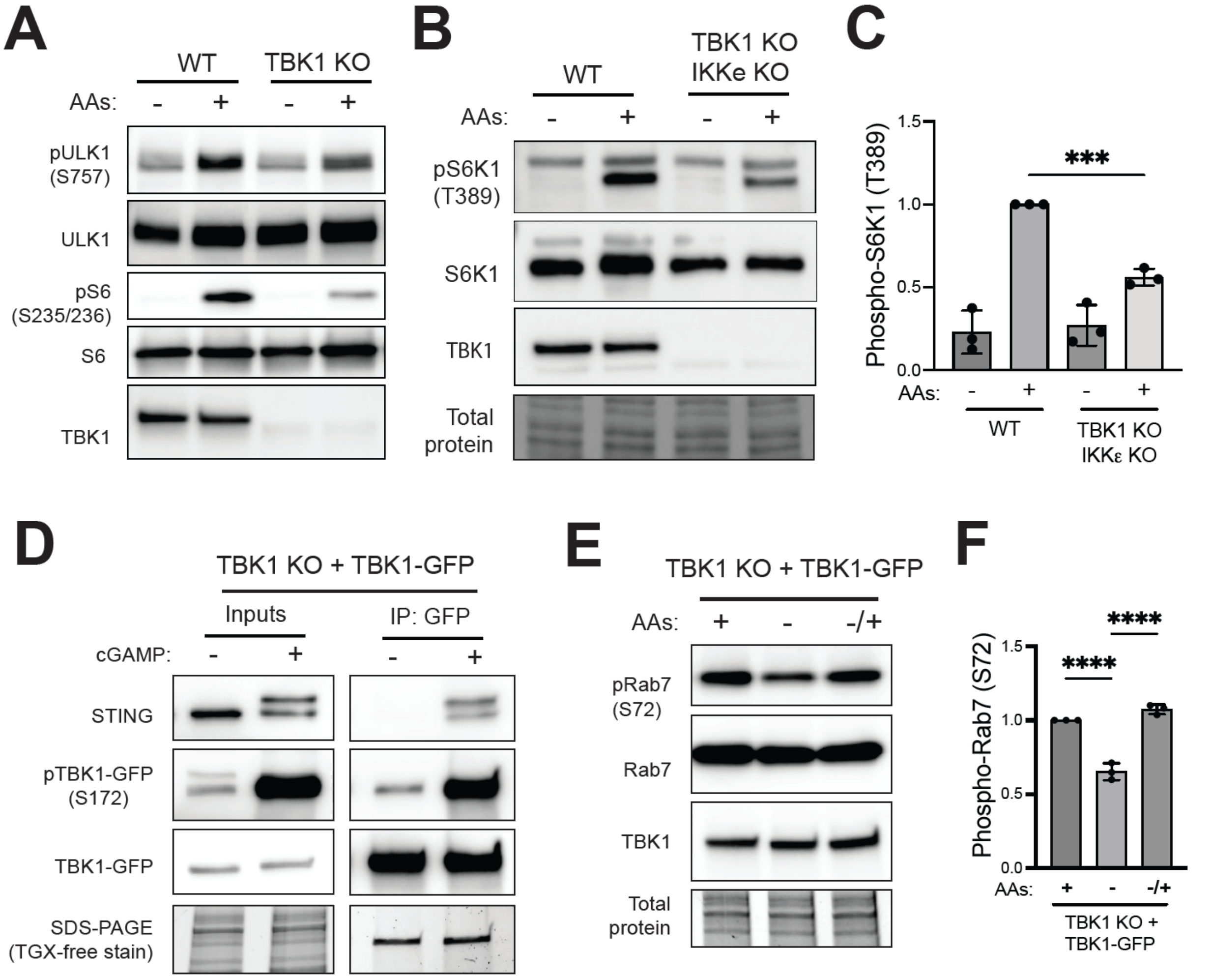


**Figure S1: TBK1 is required for efficient amino acid dependent mTORC1 and cGAMP dependent STING activation.** (A) Immunoblot analysis of phospho-ULK1 at S757, total ULK1, phospho-S6 (S235/S236) and TBK1 of WT and TBK1 KO HeLa cells starved for 60 min (-) and re-fed with amino acids (+). (B) Immunoblot analysis of S6K1 phosphorylated at Thr389 [pS6K1 (T389)], total S6K1 and TBK1 in whole-cell lysates from WT versus TBK1 KO RAW 246.7 cells that were starved of amino acids for 60 minutes (-) and then re-fed with amino acids for 60 minutes (+). TGX stain-free method was used to visualize total protein. (C) Quantification of phospho-S6K1 (T389) normalized to total S6K1 and expressed as a fold change compared to the WT cells under re-fed conditions. Statistical significance was determined by ordinary one-way analysis of variance (ANOVA) with Šidák *post hoc* test (n=3; mean ± SD; ***, $p<0.001$). (D) Immunoblot analysis of STING, phospho-TBK1-GFP and total TBK1-GFP in the lysates (Inputs) and immunoprecipitated TBK1-GFP (IP: GFP) of TBK1 KO HeLa cells rescued with TBK1-GFP untreated (-) or treated with cGAMP (70 μM) for 120 min (+). (E) Immunoblot analysis of pRab7 (S72), total Rab7 in cell lysates of TBK1 KO + TBK1-GFP HeLa cells under basal fed conditions (+), starved (-) and amino acid re-fed (-/+). (F) Phospho-Rab7 levels were quantified and normalized to total Rab7 (basal conditions were considered 1). Statistical significance was determined by ordinary one-way ANOVA with Šidák post-test (n=3; mean ± SD; ****, $p<0.0001$).

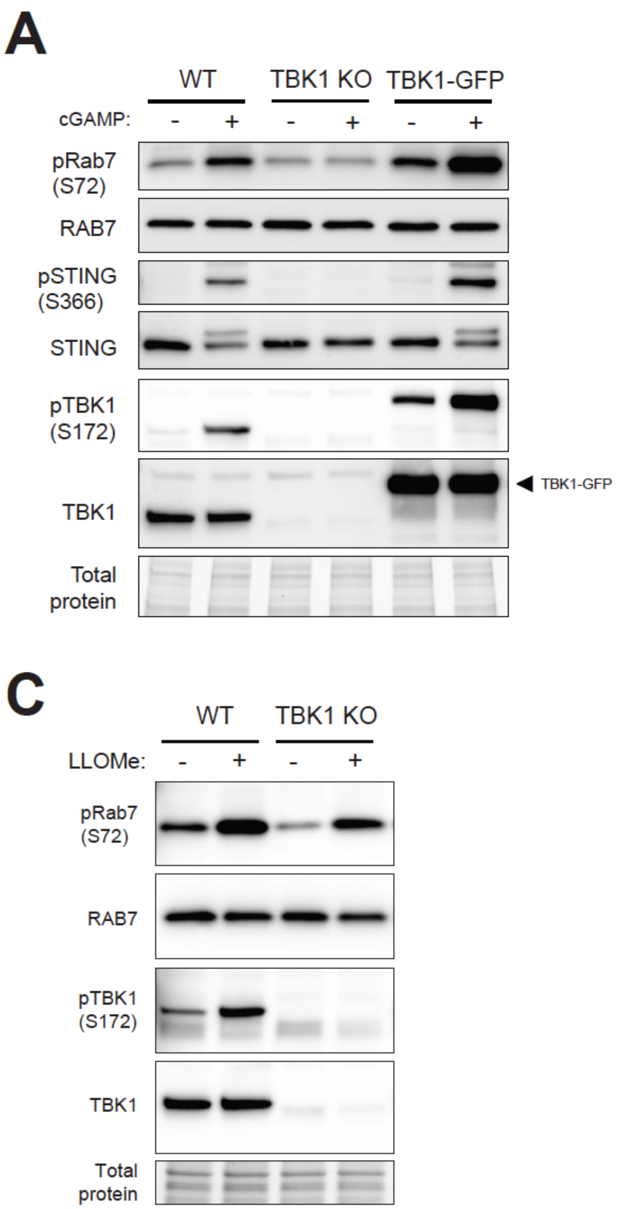


**Figure S2: TBK1 activity is regulated by multiple signals.** (A) Immunoblot analysis of the indicated proteins in WT, TBK1 KO and TBK1-GFP HeLa cells untreated (-) or treated with cGAMP (70 μM) for 120 min. (B) Immunoblot analysis of the indicated proteins of WT and STING KO RAW 246.7 cells starved for 60 min (-) and then refed with amino acids for 60’ (+). (C) Immunoblot analysis of the indicated proteins in WT and TBK1 KO HeLa cells untreated (-), DMSO 0.1 % (v/v), or treated with LLOMe (1 mM) for 30 min.

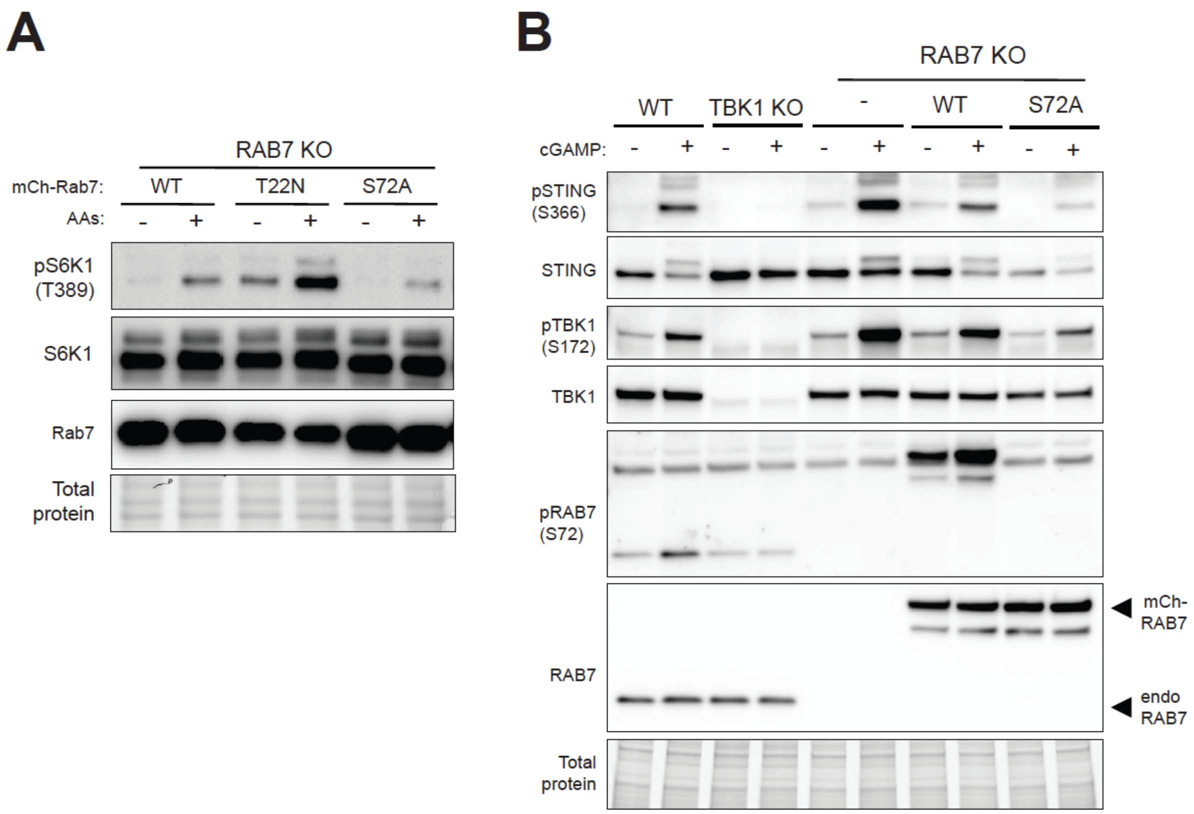


**Figure S3: Rab7 mutants show defective mTORC1 and STING signaling.** (A) Immunoblot analysis of the indicated proteins of Rab7 KO HeLa cells transiently transfected with mCherry-tagged wild-type Rab7 and mutants (T22N and S72A), starved for 60 min (-) and then refed with amino acids for 60 min (+). (B) Immunoblot analysis of the indicated proteins of WT, TBK1 KO, Rab7 KO and Rab7 KO stably expressing mCherry-tagged wild-type or S72A versions of Rab7 HeLa cells untreated (-) or treated with cGAMP (70 μM) for 120 min.

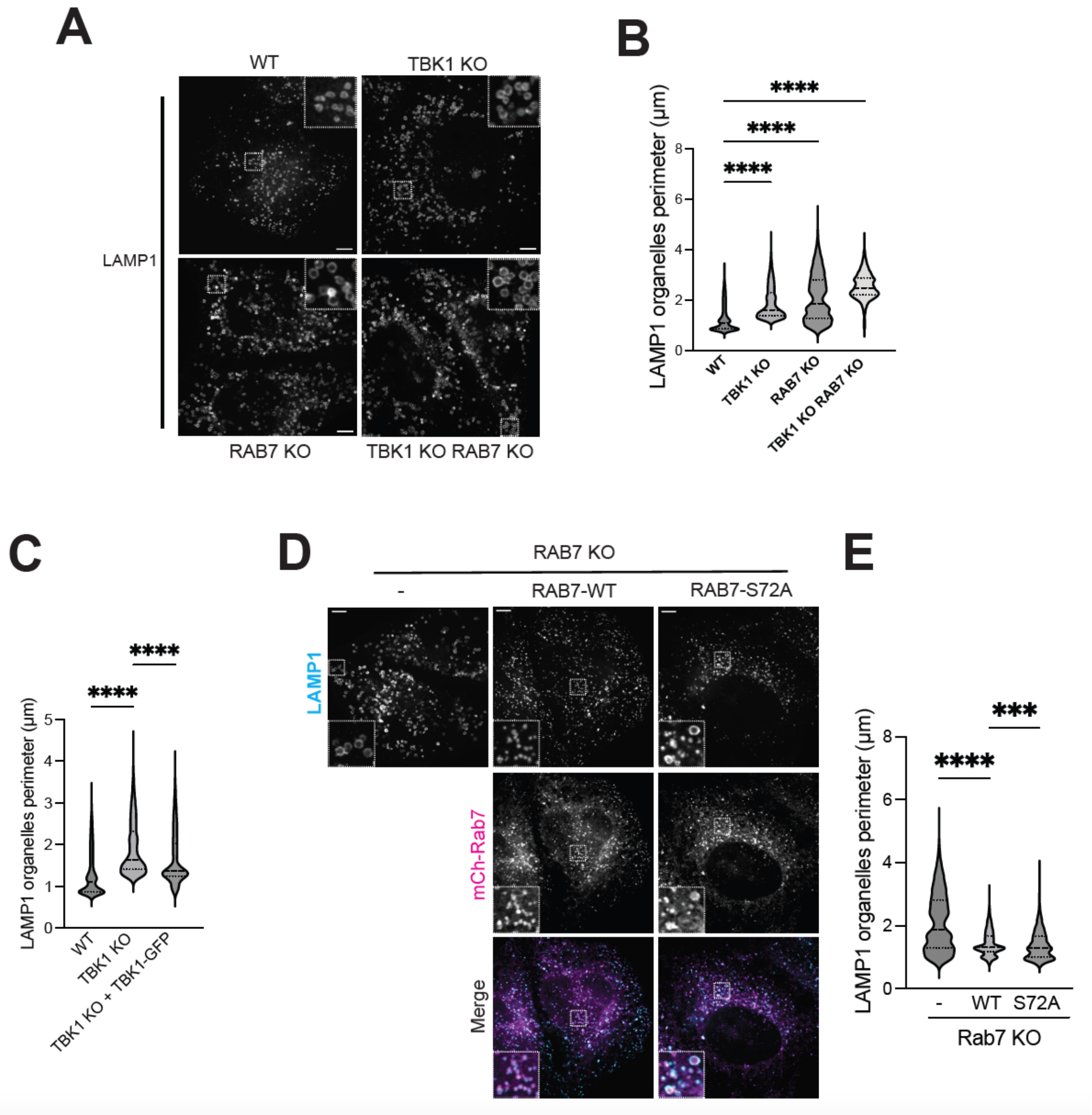


**Figure S4: TBK1 regulates lysosome size independent of Rab7.** (A) Immunofluorescence microscopy analysis of LAMP1 in basal conditions of WT, TBK1 KO, RAB7 KO, TBK1 KO RAB7 KO HeLa cells. (B) Quantification of of LAMP1-positive organelle (lysosomes) perimeter (µm) in cells of the indicated genotypes. Scale bar: 5 µm. (C) Quantification of LAMP1-positive organelle (lysosomes) perimeter from WT, TBK1 KO and TBK1-GFP rescue data presented in Figure 1G. (D) Immunofluorescence microscopy analysis of LAMP1 and mCh-Rab7 in basal conditions for Rab7 KO HeLa cells versus Rab7 KO stably expressing RAB7-WT or RAB7-S72A. (E) Quantification of LAMP1-positive organelle (lysosomes) perimeter (µm) in cells of the indicated genotypes. Data plotted in panels B, C and E represents results from 2-4 independent experiments; 11-29 regions of interest; 1-3 cells per region of interest. Statistical significance was determined by the Kruskal–Wallis's test followed by Dunn's post test (Violin Plot; ****, $p<0.0001$; ***, $p<0.001$). Scale bar: 5 μm.

### Supplemental Methods

### Supplemental Methods

**Table 1 -** Summary of cell lines used in this study.

| Cell Line | Genotype | Reference |
|---|---|---|
| HeLa M | WT | Pietro De Camilli, Yale University |
| HeLa M | TBK1 KO | This work |
| HeLa M | TBK1 KO + TBK1-GFP | This work |
| HeLa M | TBK1 KO + TBK1-E696K-GFP | This work |
| HeLa M | RAB7 KO | This work |
| HeLa M | RAB7 KO + mCherry-RAB7 | This work |
| HeLa M | RAB7 KO + mCherry-RAB7-S72A | This work |
| HeLa M | TBK1 KO + RAB7 KO | This work |
| RAW 264.7 | WT | ATCC |
| RAW 264.7 | STING KO | Bentley-DeSousa and Ferguson, 2023 |
| RAW 264.7 | TBK1 KO + IKK KO | Bentley-DeSousa and Ferguson, 2023 |

**Table 2 -** Summary of nutrients and drug treatments used in this study

| Compounds/Drugs | Company | Product Number |
|---|---|---|
| DMEM | Thermo Fisher Scientific | 11965-092 |
| HI-FBS | Thermo Fisher Scientific | 16140-071 |
| Penicillin/Streptomycin (10,000 U/mL) | Thermo Fisher Scientific | 15140122 |
| Puromycin | Thermo Fisher Scientific | A11138-03 |
| RPMI 1640 Medium Modified w/o Amino acids | USBiological | R9010-01 |
| MEM Amino Acids | Gibco | 11130-051 |
| 2’,3’-cGAMP | Chemietek | CT-CGMAP |
| BX-795 | Cayman Chemical | 14932 |
| LLOMe | Cayman Chemical | 16008 |

**Table 3** - Summary of plasmids used in this study

| Plasmid | Reference |
|---|---|
| pSpCas9-2A-Puro-TBK1-gRNA (PX459) | This paper |
| pSpCas9-2A-Puro-RAB7A-gRNA1 (PX459) | This paper |
| pSpCas9-2A-Puro-RAB7A-gRNA2 (PX459) | This paper |
| pEIF1A-piggyBac transposase | Michael Ward (NINDS) (Pantazis et al., 2022) |
| pPB-EF1A-Puro-hTBK1-EGFP | This paper |
| pPB-EF1A-Puro-hTBK1-E696K-EGFP | This paper |
| pPB-EF1A-Puro-mCherry-RAB7 | This paper |
| pPB-EF1A-Puro-mCherry-RAB7-S72A | This paper |
| pCMV-mCherry-RAB7 | Addgene #61804 |
| pCMV-mCherry-RAB7-S72A | This paper |

| | |
|---|---|
| pCMV-mCherry-RAB7-T22N | This paper |
| pCMV-myc-RAB7 | Christopher Burd, Yale University |
| pCMV-myc-RAB7-T22N | Christopher Burd, Yale University |

**Table 4** - Sequences of oligonucleotide primers used in this study

| **Primer** | **Sequence (5'-3')** |
|---|---|
| RAB7_F | CAAAAAAGCAGGCTGCCACCATGGTGAGCAAGGGCGAG |
| RAB7_R | TCAGCAACTGCAGCTTTCTG |
| PB-EF1_F | CAGAAAGCTGCAGTTGCTGAACCCAGCTTTCTTGTACAAAGTG |
| PB-EF1_R | GGTGGCAGCCTGCTTTTTTG |
| RAB7-S72A_F | ACGGTTCCAGGCTCTCGGTGT |
| RAB7-S72A_R | TCCTGTCCTGCTGTGTCC |
| RAB7-S72E_F | ACGGTTCCAGGAGCTCGGTGTGG |
| RAB7-S72E_R | TCCTGTCCTGCTGTGTCC |
| TBK1-E699K_F | ATTAAAGGAAAAGATGGAAGG |
| TBK1-E699K_R | TTCTTCATACCAAGAGTC |
| TBK1 gRNA sense | CACCGCATAAGCTTCCTTCGTCCAG |
| TBK1 gRNA antisense | AAACCTGGACGAAGGAAGCTTATGC |
| RAB7A gRNA1 sense | CACCGGTCATCCACCATCACCTCCT |
| RAB7A gRNA1 antisense | AAACAGGAGGTGATGGTGGATGACC |
| RAB7A gRNA2 sense | CACCGCATTCAAAACCCTAGATAGC |
| RAB7A gRNA2 antisense | AAACGCTATCTAGGGTTTTGAATGC |

**Table 5** - Description of antibodies used in this study

| **Antibody** | **Company** | **Concentration** |
|---|---|---|
| S6K1 | Cell Signaling Technologies | 1:2000 |
| P-S6K1 (T389) | Cell Signaling Technologies | 1:1000 |
| ULK1 | Cell Signaling Technologies | 1:2000 |
| P-ULK1 (S757) | Cell Signaling Technologies | 1:2000 |
| S6 | Cell Signaling Technologies | 1:6000 |
| P-S6 (S235/236) | Cell Signaling Technologies | 1:2000 |
| mTOR | Cell Signaling Technologies | 1:2000 |
| Rab7 (E9O7E) | Cell Signaling Technologies | 1:4000 |
| P-Rab7 (S72) | Abcam | 1:1000 |
| STING | Cell Signaling Technologies | 1:1000 |
| P-STING (S366) | Cell Signaling Technologies | 1:1000 |
| TBK1 | Cell Signaling Technologies | 1:2000, 1:500 |
| P-TBK1 (S172) | Cell Signaling Technologies | 1:1000, 1:100 |
| LAMP1 (D2D11) | Cell Signaling Technologies | 1:2000 |
| LAMP1 (H4A3) | DSHB | 1:2000 |
| GFP | Invitrogen | 1:500 |

| | | |
|---|---|---|
| PDI | Cell Signaling Technologies | 1:1000 |
| GM130 | BD Biosciences | 1:2000 |
| Rabbit IgG (HRP) | Cell Signaling Technologies | 1:2000 |
| Mouse IgG (HRP) | Cell Signaling Technologies | 1:2000 |
| Biotin (HRP) | Cell Signaling Technologies | 1:4000 |
| Alexa 488-Rabbit | Invitrogen | 1:600 |
| Alexa 568-Rabbit | Invitrogen | 1:600 |
| Alexs 647-Mouse | Invitrogen | 1:600 |
